# Dorsomedial striatal activity tracks completion of behavioral sequences

**DOI:** 10.1101/2021.04.01.437899

**Authors:** Youna Vandaele, David J Ottenheimer, Patricia H Janak

## Abstract

For proper execution of goal-directed behaviors, individuals require both a general representation of the goal and an ability to monitor their own progress toward that goal. Here, we examine how dorsomedial striatum (DMS), a region pivotal for forming associations among stimuli, actions, and outcomes, encodes the execution of goal-directed action sequences that require self-monitoring of behavior. We trained rats to complete a sequence of at least 5 consecutive lever presses (without visiting the reward port) to obtain a reward and recorded the activity of individual cells in DMS while rats performed the task. We found that the pattern of DMS activity gradually changed during the execution of the sequence, permitting accurate decoding of sequence progress from neural activity at a population level. Moreover, this sequence-related activity was blunted on trials where rats did not complete a sufficient number of presses. Overall, these data suggest a link between DMS activity and the execution of behavioral sequences that require monitoring of ongoing behavior.

## Introduction

Goal-directed behaviors depend upon outcome expectation to guide the behavioral response (Dickinson, 1989; Dickinson and Balleine, 1994). However, goal-directed responses are rarely isolated actions. They often involve the execution of complex action sequences for the goal to be attained (Dezfouli and Balleine, 2013; Dezfouli et al., 2014), for instance, in the case of a predator hunting its prey. For optimal performance in this situation, individuals not only require a general representation of the goal of their actions (catching the prey) but also need to track their own progress toward that goal (approaching the prey).

The dorsomedial striatum (DMS) plays a pivotal role in forming associations between stimuli, actions and outcomes (Balleine and O’Doherty, 2010; Balleine et al., 2009; Corbit and Janak, 2010; Yin et al., 2005). Electrophysiological recording studies typically report that neurons in DMS are prominently modulated during execution of action sequences, in response to reward-predictive cues, and at the time of reward consumption (Barnes et al., 2005; Ito and Doya, 2015; Jin and Costa, 2015; London et al., 2018; Robbe, 2018; Rueda-Orozco and Robbe, 2015; Thorn et al., 2010; Vandaele et al., 2019). Interestingly, a growing number of studies also show a role for dorsal striatum in temporal processing. Specifically, the firing dynamics of striatal populations reliably encode time over tens of seconds in Pavlovian and instrumental tasks involving delays and timing behavior (Bakhurin et al., 2017; Emmons et al., 2017; Gouvêa et al., 2015; Matell et al., 2003; Mello et al., 2015). Furthermore, inactivation of the striatum significantly impairs interval timing (Akhlaghpour et al., 2016; Emmons et al., 2017; Gouvêa et al., 2015), a cognitive process that may contribute to tracking behavioral progress toward a goal during execution of action sequences.

Given this collection of evidence that DMS is involved in the execution of behavioral sequences and timing, we set out to characterize the activity of DMS in a task requiring rats to monitor their own progress while performing lever press sequences. Specifically, rats had to complete sequences of at least 5 consecutive lever presses without entering the reward port in order to obtain a reward. Thus, some sort of monitoring of action output would improve overall reward acquisition in this task. We found that DMS activity evolved across the behavioral sequence in a ramp-like pattern of activity, permitting accurate decoding of sequence progress at the population level. Additionally, the magnitude of sequence-related activity was blunted on incomplete trials, suggesting that DMS activity may be critical for proper monitoring and execution of behavioral sequences.

## Results

### Rats monitor their performance during execution of fixed-length lever press sequences

Rats were trained to complete sequences of consecutive lever presses without checking the reward port in order to obtain a reward. Premature visits of the reward port were penalized by resetting the response requirement (Fig 1A). The acquisition of the sequence was gradual; after initial instrumental training under a continuous reinforcement schedule, the subjects (N=9) were trained with a sequence length of 2 and then 3 lever presses before training at the final sequence length of 5 lever presses (fixed sequence 5 schedule; FS5) for 16 to 24 sessions (methods). Completion of the full lever press sequence without checking the port increased over time, as indicated by the rise in percentage of complete sequences, reaching an asymptote at 80% across the last 3 recording sessions (Fig 1B; main effect of sessions: F20,140=13.96, p<0.0001), and by a decrease in the number of premature port entries (Fig 1B; main effect of sessions: F20,140=10.17, p<0.0001). We analyzed behavior and DMS neuronal spiking activity after stabilization of performance from the 8th FS5 session (Fig 1B).

**Figure 1:**
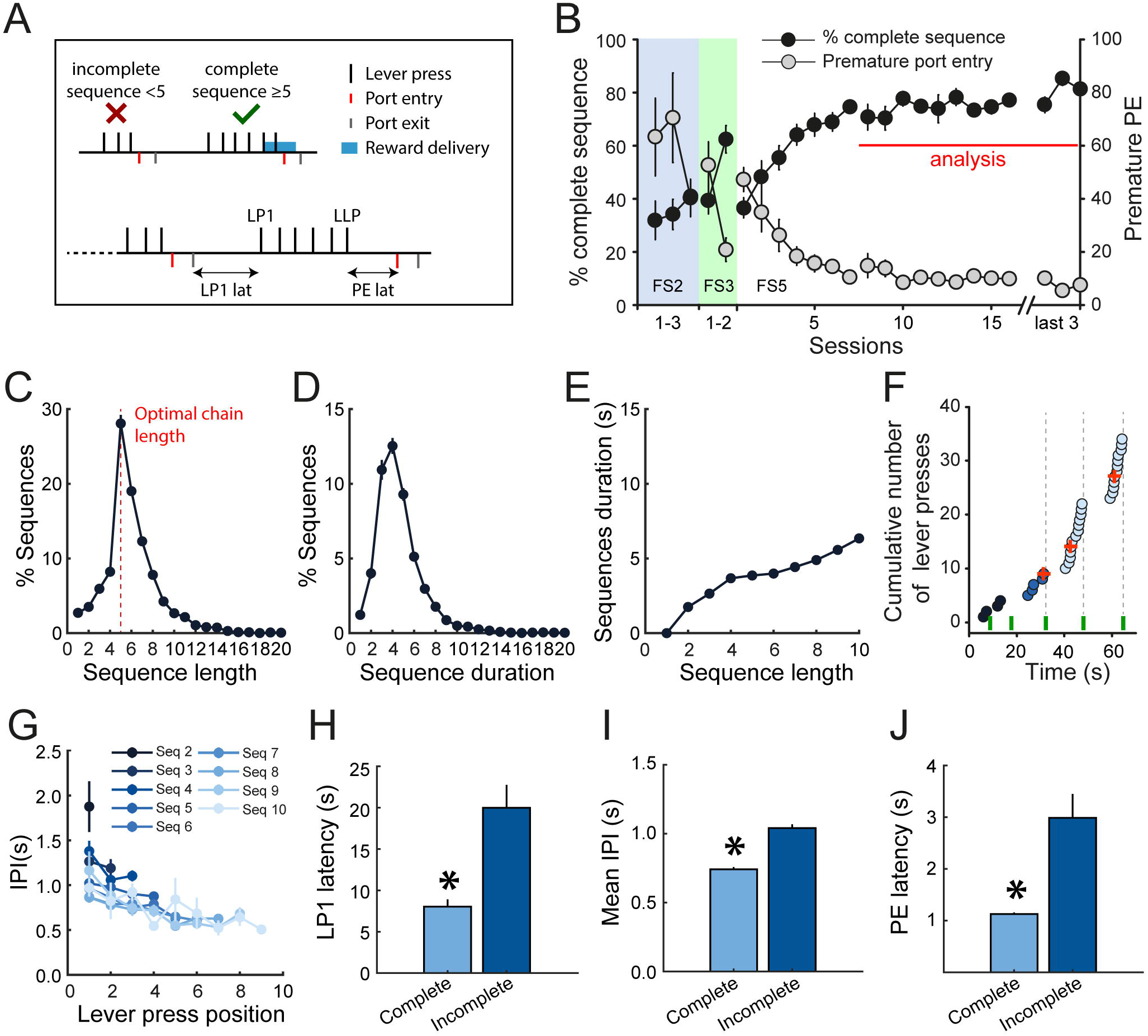
Rats monitor their performance during execution of fixed-length lever press sequences. A. Diagram of the FS5 task. LP1: first lever press. LLP: last lever press. PE: port entry. B. Mean percentage of complete sequence and premature port entry across training sessions. C-D. Mean distribution of sequence length (C) and duration (D). E. Mean sequence duration as a function of sequence length. F. Microstructure of behavior during execution of sequences in a representative rat, at the start of the session. Circles: lever presses. Green ticks: port entries. Red crosses: reward deliveries G. Mean inter-press intervals (IPI) as a function of sequence length and across lever press position. H-J. Mean LP1 latency (H), IPI (I) and PE latency (J) in complete and incomplete sequences. *p<0.001. C-E and G-J represent the mean (±SEM) of 118 FS5 sessions.

Analysis of rats’ lever pressing revealed that rats were monitoring their reward-seeking behavior to perform the task optimally (Fig 1C-D). Rats most frequently made 5 lever presses, indicating that they had learned the response contingency and titrated their behavior to the optimal chain length (Fig 1C). Rats may have also achieved high performance in the task by estimating the time elapsed from the first lever press. Analysis of the distribution of sequence durations reveals a peak at 4s (Fig 1D), which corresponds to the average time for the completion of 4 to 6 lever press sequences (Fig 1E).

We also examined whether there were any changes in pressing behavior within individual sequences that could indicate that the subjects were actively monitoring sequence progress (Fig 1F-G). Indeed, rats were faster to press the lever as they approached the end of the sequence (Fig 1F), illustrated by decreases in the mean inter-press intervals (IPI) as subjects progressed through the sequence (Fig 1G; sequences 4 to 9: F-values>4.5, p-values<0.05). Overall, rats initiated, executed, and terminated complete sequences more quickly than incomplete sequences, suggesting that they were less motivated on trials for which they did not complete the lever press requirement (Fig 1H-J; latency to the first press: F_1,116_=21.2, p<0.001; mean IPI: F_1,116_=210.9, p<0.001; PE latency: F_1,116_=16.44, p<0.001). Analyses of individual rats showed similar results (Fig S1-S4).

### DMS activity is characterized by a ramp across lever presses followed by a switch in activity at the approach of the port

To characterize the activity of putative medium spiny neurons (MSNs; N=1014, 88% of recorded units; Fig. S5) in DMS (Fig. 2A) during the execution of lever-press sequences, we normalized the time elapsed from the first (LP1) to the last (LLP) lever press (Fig. 2B; methods). Because we observed that the latency to enter the port depended on the sequence length (Fig. 1J), the same normalization approach was employed between the last lever press and the port entry (port approach period, PA).

**Figure 2:**
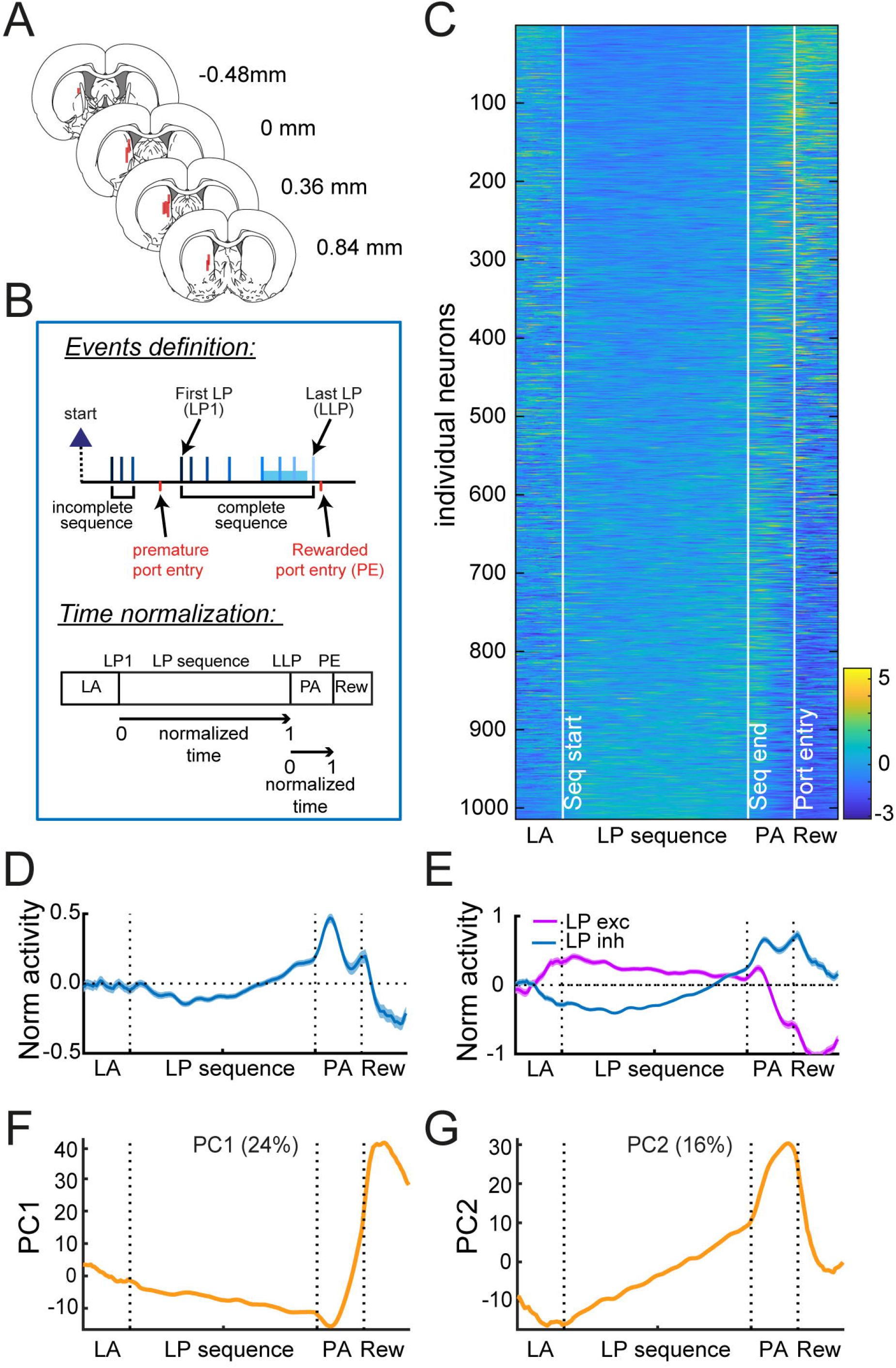
DMS activity is characterized by a ramp across lever presses followed by a switch in activity at the approach of the port. A. Electrode placements. B. Diagram of task events and normalization of spiking activity according to sequence duration and port entry latency. LA: Lever Approach; PA: Port Approach; Rew: Reward. C-D. Heatmap (C) and average z-score (±SEM) (D) of MSNs. E. Average z-score (±SEM) of MSNs excited or inhibited during lever presses. F-G Eigenvector values of PC1 (F) and PC2 (G). The % variance explained by each component is indicated.

We first analyzed MSN activity during complete sequences (greater or equal to 5 lever presses). The normalized activity of individual MSNs is depicted in Fig. 2C. While the average normalized activity of this population decreased during the first lever presses, rose toward the end of the sequence, peaked after the last lever press and dropped during the port approach (Fig 2D), the activity of individual units is clearly more variable. When neurons were separated based on the sign of their mean z-score during lever presses (excitation or inhibition), we observed a switch in activity during the port approach (Fig. 2E), with a decrease in firing for neurons excited during lever presses and an increase in firing for neurons inhibited during lever presses. We note that a comparable pattern of activity was observed when firing rate was analyzed in 40ms time bins around the lever press and port entry events (−0.25 to 0.25s peri-events; Fig. S6). We also characterized the population activity using a principal component analysis. This analysis revealed that, across the population, there tended to be ramps of activity across the sequence and sharp transitions in activity during port approach and reward acquisition (Fig. 2F-G). Similar ramps of activity were found when the PCA analysis was conducted on the lever press sequence only (Fig. S7).

### Progress in the lever press sequence is encoded in DMS activity pattern

The gradual shift in activity during sequence execution suggested that the animal’s progress in the execution of the lever press sequence could be read out from DMS activity. To test this hypothesis, we trained linear discriminant analysis (LDA) models on the normalized spike activity of individual DMS neurons across 5 equivalently-sized, consecutive intervals of the sequence on a subset of trials. We used these models to classify the sequence position of the intervals for held-out, test trials (Fig 3A). For this analysis, we pooled neurons recorded from sessions comprising at least 26 complete sequences of duration shorter than 20s and with a latency to retrieve the reward shorter than 10s (N=903). We first conducted this analysis on randomly selected pseudo-ensembles of neurons. For ensemble sizes of N=50 and above, LDA models accurately predicted the time positions of all held-out sequence intervals (Fig 3B; permutation tests: p-values<0.0001). The accuracy increased with the ensemble size (reaching 65 to 91% for N=900 neurons), and differed as a function of interval position (main effect of ensemble size, F_4,1249_=1096.7 p<0.0001; main effect of interval position, F_4,1249_=331.3, p<0.0001; interaction F_16,1249_=3.0, p<0.0001). Interestingly, the accuracy was higher at the first and last intervals compared to intermediate intervals (post-hoc p-values<0.0001). Similar results were found when LDA models were trained on activity across fewer or greater number of consecutive time intervals (Fig S8).

**Figure 3:**
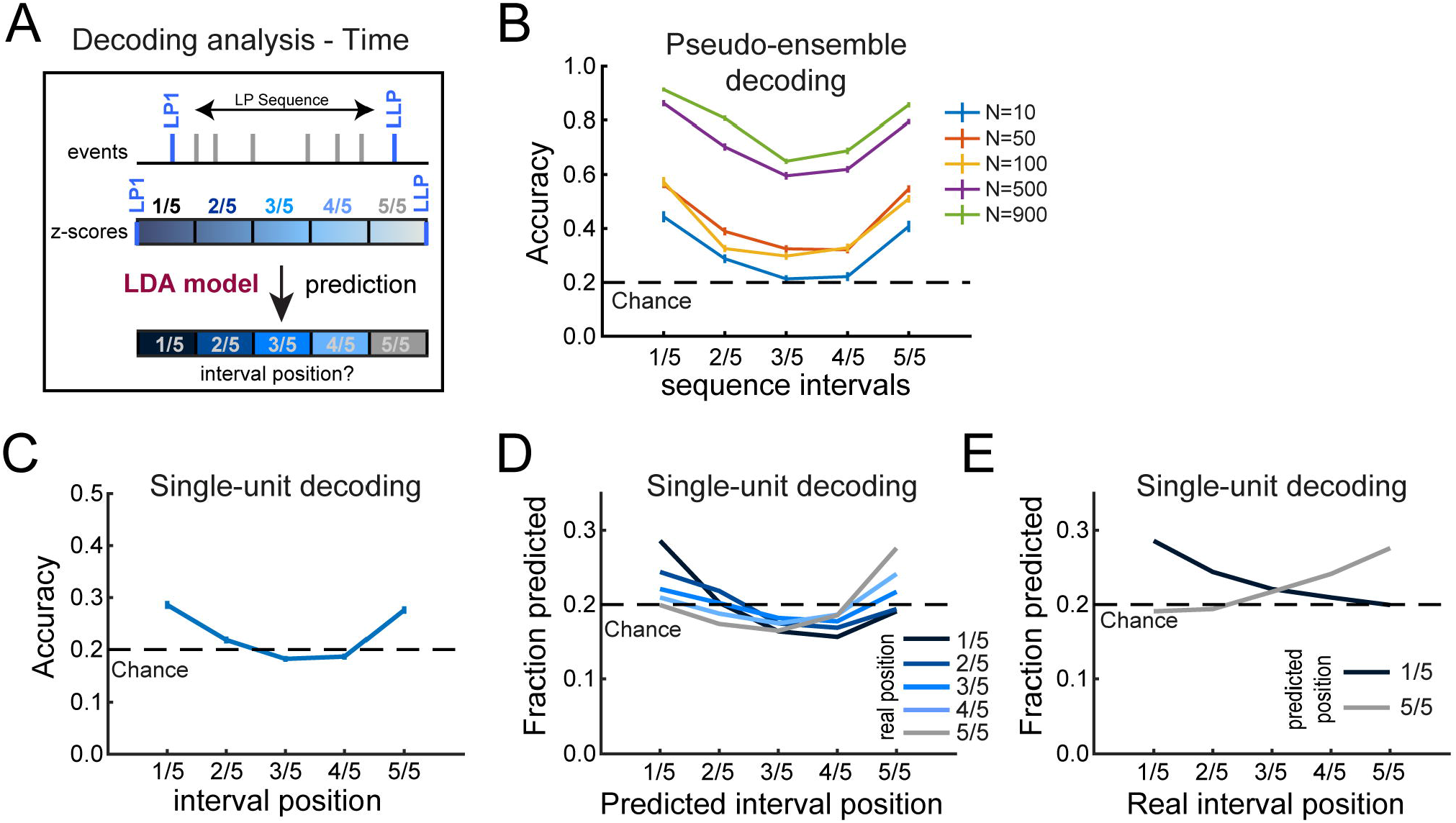
Progress in the lever press sequence is encoded in DMS activity pattern. A. Diagram of the decoding analysis. B. Mean decoding accuracy across time intervals and as a function of pseudo-ensemble size. C. Mean single-unit decoding accuracy across time intervals. D. Fraction of predicted interval position as a function of real interval position. E. Fraction of intervals predicted as the first (1/5) and the last (5/5) interval as a function of real interval position.

With evidence that sequence progress is encoded at the population level, we next investigated whether individual neurons were sufficient to decode sequence position. On average, individual neuron activity could also predict above chance the position of the first, second and last sequence intervals (Fig 3C) (permutation tests; 1^st^: p<0.0001; 2^nd^: p<0.05; last: p<0.0001), although the overall accuracy was poor. Interestingly, there was a pattern to the misclassification errors. Overall, time intervals were more likely classified as the first interval the closer they were to the sequence initiation and, correspondingly, were more likely classified as the last interval the closer they were to the sequence termination (Fig 3D). Accordingly, the fraction of neurons from which the first interval is predicted decreased (first vs last interval McNemar χ^2^=580.3, p<0.0001) whereas the fraction of neurons from which the last interval position is predicted increased (McNemar χ^2^=533.2, p<0.0001), as subjects progressed through the sequence (Fig 3E). Similar results were found by training LDA models on normalized spike activity around each individual lever press to decode the lever press position in the sequence (Fig S9). These results are consistent with the ramping pattern of activity observed in DMS neurons along the behavioral sequence (Fig. 2) and indicate that activity in DMS gradually progresses in a reliable pattern as the sequence is completed.

We next analyzed the activity of individual neurons that predicted the position of a given interval above chance by comparing single-unit decoding accuracy to the accuracy of the whole population of neurons after their activity was shuffled (Fig 4; methods). A majority of individual neurons accurately predicted at least one specific time interval (Fig 4A; p<0.01 for at least one interval in 58% of neurons, 523 out of 903), with larger proportions of good decoders for the first and last intervals (Fig 4B; 1^st^ interval N=295, 33%; 5^th^ interval: N=291, 32%). Among these, numerous neurons concurrently decoded the location of both the first and last intervals (N=140) whereas overlaps with the middle interval (3^rd^ interval) were more limited (Fig 4A; 1^st^ and 3^rd^ interval: N=53; 3^rd^ and 5^th^ interval: N=52). Since some neurons decoded several intervals’ location, we grouped neurons by their best predicted position (i.e., each neuron only represented once in this analysis), and found an effect of interval on their decoding accuracy (Fig 4C; mean for each interval: 1^st^: 0.56; 2^nd^: 0.52; 3^rd^: 0.51; 4^th^: 0.52; 5^th^: 0.53; main effect of interval: F_4,883_=5.64, p<0.001). Examination of the heatmaps (fig 4D) and the average activity pattern of best decoders (Fig 4E) illustrates that neurons decoding the first and last interval positions resembled the prominent activity patterns across the population in Fig. 2; that is, these neurons’ activity tended to ramp across the sequence, permitting reliable decoding of the beginning and end of the sequence. Interestingly, with the exception of the last interval, we systematically observed a transient inhibition during the interval best predicted by the neurons (Fig 4E). Similar activity patterns were observed when the LDA models were trained on the normalized spike activity around each individual lever press (Fig S10). This analysis demonstrates that, at the level of individual neurons, there is robust encoding of aspects of the sequence, characterized by a ramp toward termination of the sequence, and single-unit inhibition during the prior predicted intervals. Yet, none of the neurons encoded every interval of the sequence, suggesting that strong population encoding of behavioral progress toward a goal (Fig 3B) emerges from single units individually encoding fewer time intervals but collectively encoding the full behavioral sequence at a population level.

**Figure 4:**
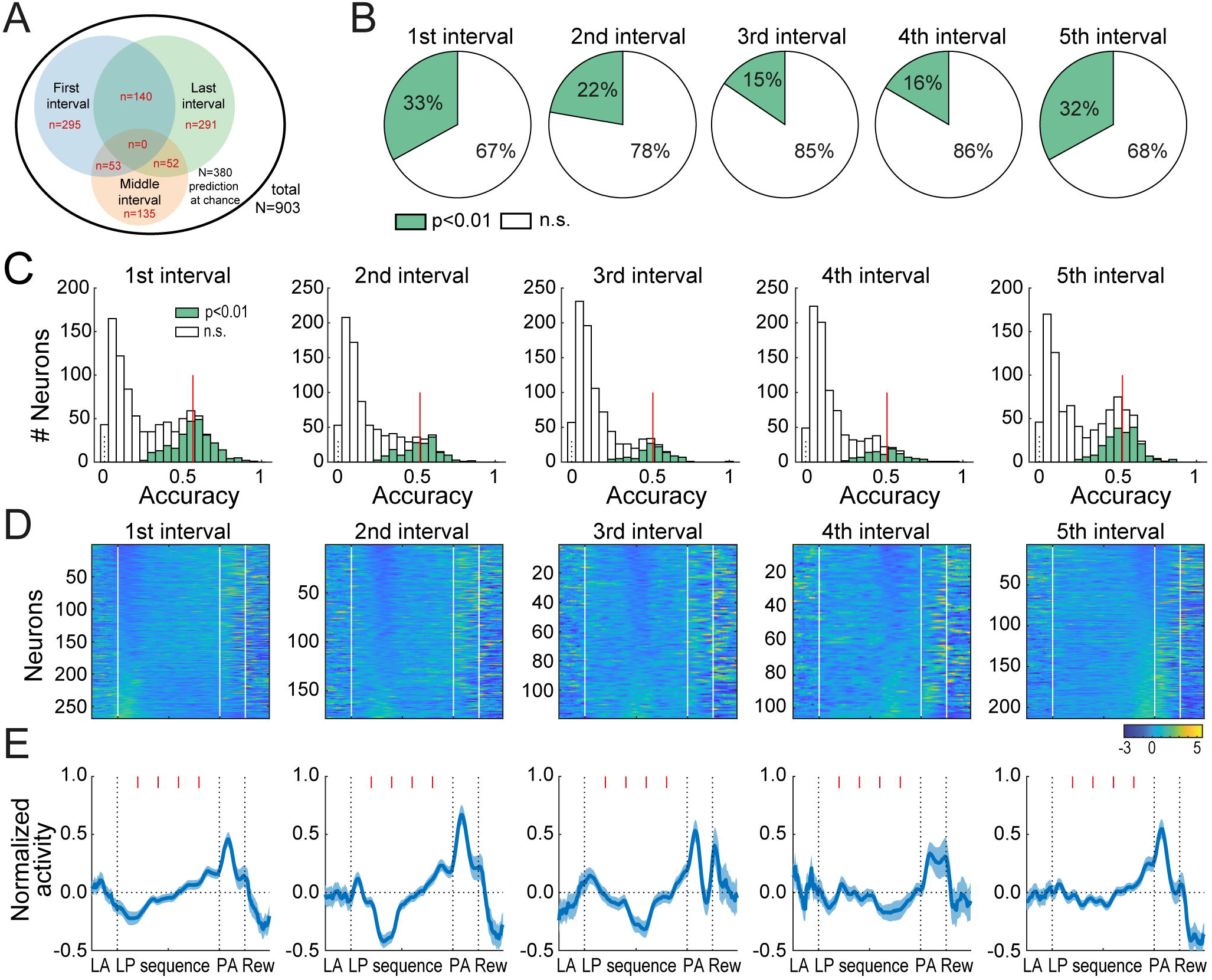
Stronger decoding at beginning and end of sequence by individual neurons. A. Venn diagram of neurons predicting the position of the first, third and last interval above chance. B. Proportion of individual neurons that best predicted the position of each intervals above chance, beginning at the first lever press and ending at the last lever press. C. Distribution of decoding accuracy of individual neurons that best predicted each of five time intervals. D-E. Heatmaps (D) and average z-score (±SEM) (E) of neurons that best predicted the position of an interval above chance, for each interval. Red ticks in E mark the limit of consecutive intervals. Activity during the lever approach, port approach and reward consumption is shown but not included in the decoding analysis.

### Attenuated activity pattern during incomplete sequences

The sequence decoding analysis demonstrates that DMS activity tracks rats’ progress in the execution of complete lever press sequences. We next sought to determine whether DMS activity differs when rats fail to execute a complete sequence, and, if so, when that difference might emerge. We trained LDA models on normalized spike activity during specific time points across complete and incomplete sequences to classify held-out trials as complete or incomplete (Fig 5A). For this analysis, we pooled neurons from sessions comprising at least 10 complete and 10 incomplete sequences (N=164). We made no assumption on whether DMS activity could differentiate complete versus incomplete sequences from neural activity prior to, after, or during performance of the lever presses themselves. Thus, we separately assessed the decoding accuracy from independent analyses of the activity at the beginning and end of the sequence (0 to 0.5s post-LP1 and −0.5 to 0s pre-LLP), but also outside of the sequence during the lever approach (−1 to 0s pre-LP1), the port approach (from LLP to PE), and the time of expected reward consumption (0 to 1s post-PE) (Fig 5A). We assessed whether the decoding accuracy significantly departed from chance by comparing it with the accuracy after shuffling complete and incomplete sequences.

**Figure 5:**
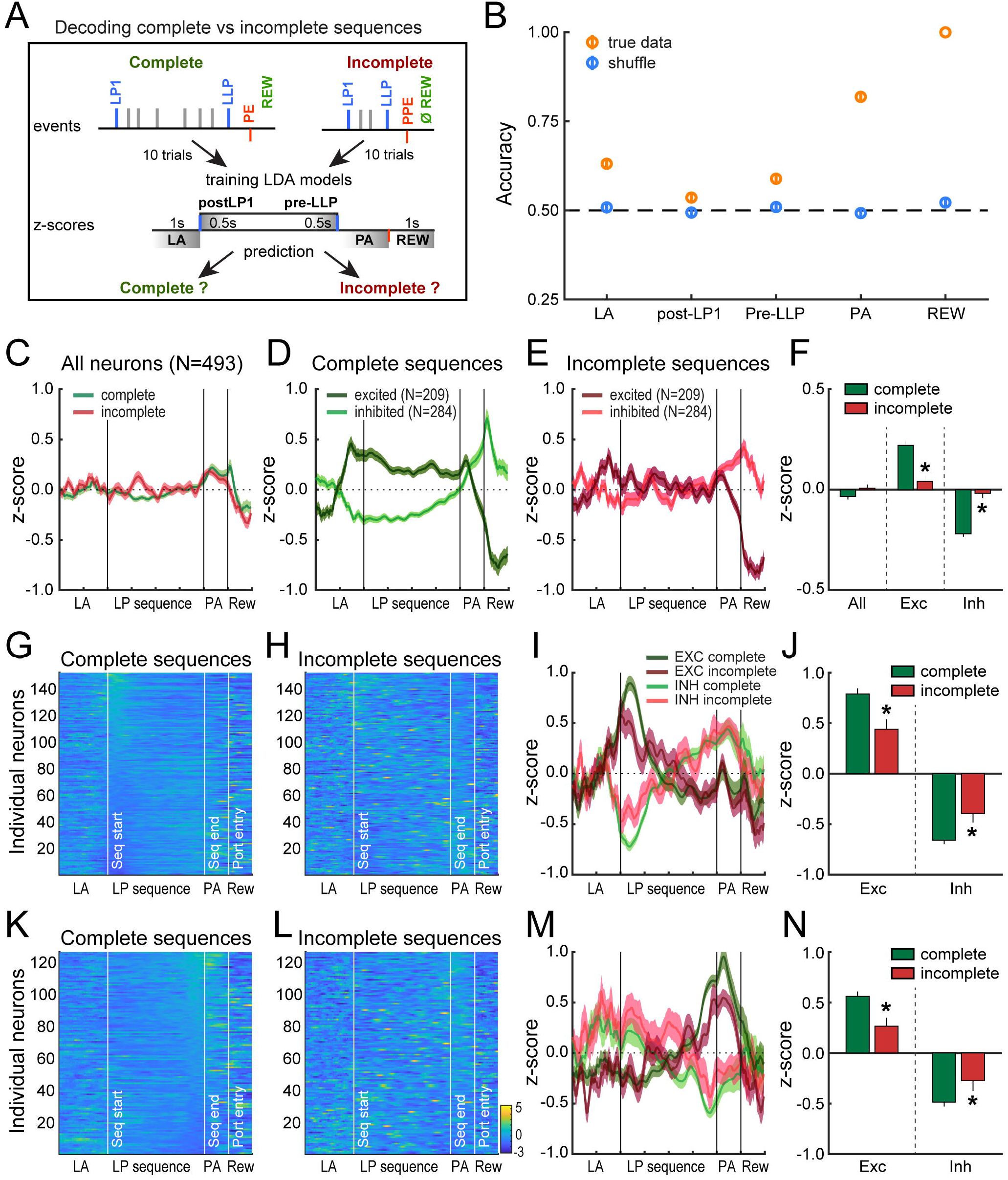
Attenuated activity pattern during incomplete sequences. A. Diagram of the decoding analysis. B. Decoding accuracy in true and shuffled conditions across time events in the behavioral sequence. Independent LDA analyses were conducted separately for each time event. C. Average z-score (±SEM) of MSN in complete and incomplete sequences. D-E. Mean z-score (±SEM) of MSN in complete (D) and incomplete (E) sequences, separated based on their mean activity during lever presses in complete sequences. F. Mean z-score (±SEM) during lever presses in complete versus incomplete sequences for all MSN and for excited or inhibited MSN. *p<0.0001. G-H. Heatmaps of first interval decoders during complete (G) and incomplete sequences (H). I-J. Average z-score (±SEM) of first interval decoders, separated based on their modulation during the first interval in complete versus incomplete sequences. J. Average z-score (±SEM) of excited and inhibited first interval decoders during the first interval. *p<0.01. K-N. Same as G-J for the last interval decoders. *p<0.05.

LDA models accurately distinguished complete from incomplete sequences at each time point of the sequence, but there were significant differences across events (F_4,499_=370.0, p<0.0001). Specifically, mean accuracy for assignment to complete or incomplete sequences was significantly above chance for all events (Fig 5B and S11; permutation tests: p-values<0.0001, except that p<0.05 for the beginning of the sequence). Complete and incomplete sequences could be dissociated from normalized DMS activity before the first lever press when rats approached the lever to initiate the sequence (mean accuracy=0.63±0.01), suggesting that a difference in neural state preceded the initiation of complete versus incomplete sequences and was maintained throughout their execution and termination. This difference in neural state parallels well the difference in motivational state illustrated by longer first lever press latency, mean IPI and port entry latency in incomplete sequences relative to complete sequences, showing that rats were less engaged in the task on trials when they made a premature port entry (Fig 1H-J). We found higher decoding accuracy at the port approach (Fig 5B, mean accuracy=0.82±0.01) and, as would be expected, perfect accuracy at the time of reward feedback (mean accuracy=1). The sensory cues (sight, smell, sound of reward delivery) that signal reward delivery in complete trials, and its absence in incomplete trials, might contribute to the relatively high decoding accuracy at the port approach time point. Surprisingly, the accuracy to classify sequences as complete or incomplete was the lowest at the beginning and termination of the sequence (Fig 5B, mean accuracy, post-LP1:0.54±0.01, pre-LLP: 0.59±0.01), suggesting that task accuracy is better predicted from the neural activity outside of the sequence. Importantly, with the exception of the first trial, incomplete sequences were evenly distributed throughout recording sessions (Fig S12).

Since we were able to dissociate complete and incomplete sequences based on DMS activity, we next asked how DMS activity differed between these two types of trials. We selected MSNs from sessions comprising at least 5 complete and incomplete sequences (N=493). Comparison of the average z-score along the lever press sequence revealed no obvious difference between complete and incomplete sequences (Fig. 5C and F; t=−1.85, p=0.065). However, when neurons were separated based on the sign of their mean z-score during lever presses in complete sequences (excitation N=209 or inhibition N=284), we observed stronger modulations in activity during execution of compete sequences (Fig. 5D) compared to incomplete sequences (Fig. 5E), with larger z-scores during execution of the sequence (Fig 5F; excitation t=5.7, p<0.0001; inhibition t=−7.91, p<0.0001). In contrast, during incomplete sequences, we observed a flatter pattern during lever approach and lever presses and a lower peak during the port approach in inhibited neurons (Fig 5E). These results suggest that the activity pattern of DMS neurons is attenuated when rats are less engaged in the task during incomplete sequences.

We further examined the activity of neurons previously identified as encoding the first and last intervals of lever press sequences (Fig 4) on complete and incomplete sequences (Fig 5G-H, K-L; 1^st^ interval predictors N=153; last interval predictors N=127 in these sessions). When neurons encoding the first time interval of complete sequences were separated according to the direction of their modulation during the first interval, we observed a larger peak in activity for complete sequences relative to incomplete sequences (Fig 5I-J; first interval excitation: t=3.72, p<0.001; inhibition: t=−3.2, p<0.01). When neurons encoding the last interval of complete sequences were separated according to their modulation at the end of the sequence, we also observed a larger peak in complete sequences compared to incomplete sequences, specifically in neurons increasing their firing rate (Fig 5M-N; last interval excitation: t=3.14, p<0.01; inhibition: t=−2.05, p<0.05). These results show that disengagement from the task during incomplete sequences is accompanied by dampened activity pattern in DMS neurons encoding behavioral progress at the initiation and termination of the lever press sequence.

## Discussion

Here, we sought to determine if on-line tracking of the execution of behavioral sequences could be observed in DMS spiking activity. Using a task in which subjects were penalized for checking for reward prior to sequence completion, we found that rats learned to track their own behavior and titrated their responding to the optimal response chain length. Efficient performance in this task was associated with a specific activity pattern in the DMS, characterized by a ramp across lever presses followed by a switch in activity as rats approached the port to retrieve the reward. It was possible to predict from this activity pattern 1) progress in the execution of the sequence and 2) whether a sequence was complete or not. Together, these results suggest that neural activity in the DMS tracks progress in the execution of action sequences, and may allow rats to time when to check the port to retrieve the expected reward.

Rats carefully monitored execution of lever press sequences in the FS5 task, most often making five consecutive responses, corresponding to the optimal chain length for the maximization of reinforcement rate. This pattern of responding is consistent with other tasks requiring execution of action chains with a minimal number of lever presses for the reward to be delivered (Rivalan et al., 2007; Vandaele et al., 2018). In absence of feedback cues indicating the correct completion of the sequence, rats had to rely on a representation of their progress in the sequence to determine when to check the port, either by estimating the number of lever presses emitted or by estimating the time elapsed from the first lever press. Interestingly, we observed an acceleration of lever press responses as rats progressed in the execution of the sequence, a “scalloping pattern” also found in fixed interval schedules involving timing behavior across delays (Dews, 1978; Wearden and Lejeune, 2006). This response pattern is thought to represent an increase in expectation as rats get closer to the expected time of reward delivery and suggests that efficient performance in the FS5 task may involve time processing.

The ramping pattern of neural activity found here is also reported in studies investigating striatal encoding of time during Pavlovian and instrumental tasks involving delays and timing behavior (Donnelly et al., 2015; Emmons et al., 2017; Gouvêa et al., 2015; Matell et al., 2003; Mello et al., 2015). In the present study, DMS spiking activity was characterized by a smooth ramp across lever presses followed by a switch in activity as rats approached the port to retrieve the reward. Upward and downward ramps across lever presses could be an expectation signal that grows as rats progress closer to the reward (Apicella et al., 1992; Hikosaka et al., 1989; Howe et al., 2013) whereas the switch in activity at the port approach could represent a reorganization of striatal ensembles as rats terminate the lever press sequence to select the port entry action (Graybiel and Grafton, 2015; Mink, 1996). The ramps reported here could have resulted from the time normalization process we used, leading to a misalignment of lever press events combined with an increase in response rate toward the end of the sequence (Fig 1). However, a similar ramping pattern was observed when we used an event-centered approach, ensuring the times of each lever press were aligned across trials (Fig S6). Although press-related peaks were observed with this approach, the peaks were higher later in the sequence, suggesting that action-related activity cannot account alone for the ramping pattern. This result suggests that DMS spiking activity integrates information about time and actions, as previously suggested (Emmons et al., 2017; Mello et al., 2015). Our results are however inconsistent with another study reporting action-related activity but no ramping pattern in the dorsal striatum during execution of lever press sequences (Ma et al., 2014a). Differences in task requirement may explain this discrepancy. While our task required rats to estimate their own progress in the sequence to reach the port at the right time, rats in the study of Ma et al had to learn the correct sequential order of lever press responses but were not required to estimate the time elapsed or the number of lever presses emitted. This suggests that striatal ramping pattern during an action sequence is only observed when animals are actively tracking relative time or progress in its execution.

Previous findings indicate that ramping activity patterns in the striatum allow time estimation and that these ramps are scalable across multiple delays (Bakhurin et al., 2017; Emmons et al., 2017; Gouvêa et al., 2015; Mello et al., 2015). In agreement with these studies, we could predict progress through sequence execution from DMS neural activity with high accuracy for larger pseudo-ensembles of neurons. The higher decoding accuracy at the start and end of the sequence is striking and suggests stronger striatal encoding of the beginning and end of the sequence compared to its intermediate parts, perhaps because the first lever press “resets” an internal clock and triggers the ramp onset whereas activity at the end of the sequence reaches a threshold for the selection of the next port entry response (Matell and Meck, 2000; Narayanan, 2016). Interestingly, several studies have shown the emergence of neuronal excitations in the dorsal striatum at the borders of behavioral sequences, as individual actions are chunked into behavioral units across sequence learning (Jin and Costa, 2010; Jin et al., 2014; Jog et al., 1999; Martiros et al., 2018; Smith and Graybiel, 2016). Although the proposed notions of task-bracketing activity or start/stop-related activity remains a matter of debate (Robbe, 2018; Sales-carbonell et al., 2018; Vandaele et al., 2019), the stronger decoding accuracy at the beginning and end of the sequence may suggest a striatal encoding of sequence initiation and termination. While changes in spiking activity were observed at the termination of the sequence during the port approach, the approach of the lever at sequence self-initiation was not characterized by phasic excitation or inhibition. More research is thus needed to directly relate our results to the presence of discrete start/stop signals in the dorsal striatum. Although the decoding accuracy was high for the full lever press sequence in large pseudo-ensembles of neurons, at the level of individual neurons, it was possible to predict the position of one or two time intervals with a moderate accuracy. Our findings therefore suggest that at a population level, DMS neurons collectively track progress in the lever press sequence by integrating single-unit encoding of fewer time intervals.

Complete and incomplete sequences could be distinguished based on DMS spiking activity at several phases of the behavioral sequence. As expected, incomplete sequences were perfectly predicted from DMS activity after the port entry, when rats realized that the reward was not delivered. The decoding accuracy significantly departed from chance during execution of the sequence and reached 80% during the port approach, when reward expectation was the highest. However, we cannot exclude that sensory cues gradually predicted reward availability in complete trials, and its absence in incomplete trials, as rats approached the reward port. Paradoxically, while the beginning and end of the sequence could be predicted from neurons’ activity with a great accuracy, it was difficult to predict from these time points whether a sequence would be completed or not. Indeed, the mean decoding accuracy was only 0.54 and 0.59 after the first and before the last lever press, respectively. These results suggest that spiking activity did not substantially differ at the beginning and end of complete versus incomplete sequences and may constitute an indirect indication that ramping pattern of striatal activity instead scales with the sequence length. It is however worth noting that the decoding analysis was conducted on a limited number of trials (10 of each) and neurons (N=164) due to the low number of incomplete sequences. This limitation may contribute to the overall lower predictive accuracy in this analysis. Future research at earlier stages of training, when rats still make a high number of incomplete sequences, would thus be desirable, and as well, could reveal how striatal activity during complete versus incomplete sequences changes in step with behavior, as rats learn to monitor their own behavior.

Surprisingly, we also found significant decoding accuracy in the classification of complete versus incomplete sequence when analyzing spiking activity during the approach of the lever. In other words, it was possible to predict whether or not a sequence would be completed before it was initiated. This result indicates that the neural activity differs between complete and incomplete sequences before sequence execution, which may parallel a difference in motivational state. Indeed, longer sequences were initiated, executed and terminated faster than short sequences, suggesting transient changes in motivation within the course of the session, rats being less motivated or less engaged in the task during shorter, incomplete sequences. This task disengagement was associated with attenuated activity patterns during incomplete sequences compared to complete sequences. Furthermore, neurons predicting the location of the first and last intervals were also less modulated at these times during incomplete sequences. This dampened activity pattern during incomplete sequence may constitute a neural marker of spontaneous failure to track execution of the sequence. However, since complete and incomplete sequences could be dissociated from activity at the lever approach and before initiation of the lever press sequence, this hypothesis is insufficient. Instead, the emission of a premature port entry could result from incorrect planning and motor impulsivity (Dalley and Robbins, 2017; Dalley et al., 2011). However, unlike incomplete sequences, impulsive actions are associated with faster response latencies than actions involving planning (Dalley and Robbins, 2017; Dalley et al., 2011), and impulsive actions in the 5-choice serial reaction time task are not associated with attenuated striatal activity patterns (Donnelly et al., 2015). Therefore, we suggest that the attenuated activity pattern in DMS, in which there is a lower modulation of firing rate during execution and at sequence boundaries, could represent a lower motivational state of the animal, resulting in premature cessation of the sequence.

To conclude, we have shown that DMS neurons encode progress toward a goal during execution of action sequences when animals are required to track their own behavior for efficient performance. This striatal region receives numerous inputs from cortical areas, notably the prefrontal cortex (Hart et al., 2018; McGeorge and Faull, 1989). Ramping patterns of activity have been found in prefrontal brain regions and are proposed to play a role in top-down control of time processing in the dorsal striatum (Donnelly et al., 2015; Emmons et al., 2017; Kim et al., 2013; Ma et al., 2014b, 2014a; Narayanan and Laubach, 2006; Parker et al., 2014; Xu et al., 2014). In addition, a ramping pattern of dopamine release emerges in the striatum as rats move toward distant goals (Howe et al., 2013) or during execution of lever press sequences (Collins et al., 2016; Wassum et al., 2012). This tonic ramp in dopamine signaling observed as subjects traverse real or virtual distance is proposed to reflect reward expectation (Howe et al., 2013) or the instantaneous reward prediction error (Kim et al., 2020), and may serve to support ongoing motivation to respond through a task. The importance of dopaminergic systems on timing behavior and motivational control is well reflected by the severe impairments observed in patients suffering from Parkinson’s disease (Parker et al., 2013). Further research is needed to determine how the dorsomedial striatum integrates time estimates from prefrontal regions and motivational signals from dopamine circuits in tracking progress toward a goal during execution of action sequences.

## Materials and methods

### Subjects

Long Evan rats (N=9, 4 males, 5 females; Envigo) were used in this experiment and trained during the light cycle of the temperature (21° C) and light-controlled vivarium (12-h light-dark cycle, lights ON at 7am). Rats were individually housed and maintained under light food restriction (90% of free feeding weight, water *ad libitum*). This study was carried out in accordance with the recommendations of the Guide for the Care and Use of Laboratory Animals (Institute of Instrumental Training Laboratory Animal Resources, Commission of Life Sciences, National Research Council, 1996). The protocol was approved by the institutional animal care and use committee of Johns Hopkins University.

### Behavioral training

Rats were first trained to retrieve small aliquots of a solution of 20% sucrose (0.1mL delivered over 3s) during a single magazine training session (random interval 60s for 30 minutes). Rats were then trained under a fixed ratio 1 schedule of reinforcement for 3 to 5 sessions (session limits: 1h or 30 reward deliveries). After acquisition of instrumental responding, rats underwent surgeries and training in the fixed sequence 5 task began following the post-operative period of 5 days.

In this task, rats had to repress entering into the port before the completion of sequences of at least 5 consecutive lever presses to obtain a reward, whose delivery was not signaled. Premature port entries (before completion of the ratio) were penalized by resetting the ratio, and the preceding lever press sequence was considered as incomplete. Additional presses after completion of the ratio were without consequences and considered as part of the complete sequence. Rats were first trained with a fixed sequence length of 2 lever presses for a minimum of 3 sessions or until they earned 30 rewards. The response requirement was then increased to 3 lever presses for a minimum of 2 sessions (or until earning 30 rewards) before training in the final fixed sequence length 5 schedule (FS5) for 16 to 24 sessions. Sessions were limited to 30min or 30 reward deliveries. We only analyzed neuronal activity during FS5 sessions after stabilization of performance in the current study (from the 8^th^ FS5 session, for 9 to 17 sessions). The house-light, located on the ceiling of the operant chamber, housed within sound-attenuating boxes (Med Associates, St Albans, VT) remained illuminated during sessions.

### Surgeries and recording

Rats were implanted with unilateral arrays of 8 electrodes (0.004’ insulated tungsten, with 2 silver ground wires) aimed at DMS with the following coordinates: +0.25 mm AP, ±2.3 mm ML, −4.6 mm DV. Surgeries were performed under isoflurane anesthesia (0.5-5%) with application of topical lidocaine for local analgesia and pre-operative injections of antibiotic (cefazolin: 75mg/kg) and analgesic (carprofen: 5mg/kg). Rats were first accustomed for a few sessions under FR1 to tethering with the recording cable, before training in the FS5 task. Cables connecting rats’ headsets to a commutator allow free movement throughout acquisition of single-unit activity with the multichannel acquisition processor (MAP) neural recording system (Plexon Inc, TX). Electrode arrays were lowered at the end of every other session by 160 µm increments with a microdrive. To avoid duplicates, only units from one session were included for the analysis at any electrode location.

### Analysis of electrophysiological recordings

#### Spike sorting

The multichannel acquisition processor (MAP) neural recording system (Plexon Inc, TX) was used to store and process amplified signals and timestamps of behavioral events. Analyses of interspike intervals distribution, auto-correlograms and cross-correlograms were conducted using Offline Sorter v3 and Neuro-Explorer 3.0 (Plexon Inc, TX) to isolate individual unit offline. Timestamps and waveforms were exported from Neuro-Explorer 3.0 to Matlab (MathWorks, MA) for further analysis. Analyses were restricted to units with well-defined waveforms and constant characteristics throughout the entire recording session.

#### Waveform analysis

All the analyses in this study were restricted to neurons classified as putative Medium Spiny Neurons (MSN) according to waveform and firing rate properties. Units classified as putative interneurons were excluded. Putative fast spiking interneurons (FSI) were defined by a firing rate higher than 20Hz and narrow waveforms with half-valley width lower than 0.15ms (N=41; 3.6%). Units were classified as tonically active interneurons (TAN) when the firing rate was lower than 5 Hz and the half-valley width was higher than 0.45ms (N=17; 1.5%). Neurons not classified as interneurons but showing features intermediate to MSNs and interneurons were also excluded (range 12.5-20Hz in firing rate and 0.4-0.45ms in half-valley width; N=80). As previously reported, population of putative-FSI and –TAN represented less than 5 and 1% of recorded units, respectively (Martiros et al., 2018; Schmitzer-Torbert and Redish, 2008; Stalnaker et al., 2016) (Fig S5).

#### Definition of task events and normalization of sequence related activity

Complete sequences were defined by sequences of at least 5 lever presses preceding a rewarded port entry whereas incomplete sequences comprised less than 5 lever presses and terminated with a premature port entry. Sequences longer than 20s or followed by a latency to enter the port longer than 10s were not considered. Activity during complete and incomplete lever press sequences was normalized according to the sequence duration. Specifically, the time to each spike in sequences was divided by the sequence duration, such that the first and last lever presses were considered as time 0 and 1 respectively. 100-bin histograms were generated for each sequence and frequency values were divided by the bin duration to estimate the firing rate. Similarly, activity during the port approach was normalized according to the port entry latency, such that the last lever press and the port entry were considered as time 0 and 1, respectively. 25-bins histograms were generated for each port approach and frequency values were divided by the bin duration to estimate the firing rate. The average bin width was 49.5±0.4ms for the lever press sequence (100 bins) and 41.5±0. 3ms for the port approach (25 bins). The 1s periods preceding the first lever press and following the port entry were included in analysis and corresponded to the lever approach and reward consumption, respectively. Activity during these periods was computed using 25 time bins of 40ms. Concatenated firing rate of individual neurons during lever approach, lever press sequence, port approach and reward consumption periods (consisting in a 175-bins vector) was smoothed (Matlab function makedist, half-normal distribution, mu=0, sigma=5) and z-scored as follow: (F_i_-F_mean_)/F_sd_. F_sd_ and F_mean_ represent the standard deviation and mean firing rate across the full behavioral sequence, and F_i_ is the firing rate at the i^th^ bin of the behavioral sequence. Individual neurons were considered as excited or inhibited during lever pressing if the average z-score from the first to the last lever press was positive or negative, respectively.

The same analyses were conducted using an event-centered approach (fig S6). Firing rate was analyzed during 0.4s time windows around each of the 5 lever press events and around the port-entry (40ms time bins, 20 bins per event). The lever press events were defined as follow: the 1^st^ and 2^nd^ lever presses, one randomly selected intermediate lever press, the second to last lever press and the last lever press (Fig S6). Firing rate across the behavioral sequence (consisting in a 160 bins vector) was smoothed and z-scored as described above.

#### Principal component analysis (PCA)

A principal component analysis (pca function in MATLAB) was conducted on the activity during the full behavioral sequence, including lever approach, lever presses, port approach and reward consumption periods. Spiking activity during lever presses was normalized according to the sequence duration, as described above. Similarly, activity during the port approach was normalized according to the port entry latency. This analysis was restricted to complete sequences and was conducted on a matrix of 1014 variables (number of DMS neurons included in the analysis) and 175 observations (number of time bins). The score values of the first two principal components (PCs) representing the most prevalent activity patterns among the neuronal population were analyzed. A second principal component analysis was conducted on the normalized activity restricted to the lever press sequence (Figure S7). The matrix for this analysis consisted in 1014 variables (number of DMS neurons included in the analysis) and 100 observations (number of time bins during the lever press sequence)

#### Decoding

A linear discriminant analysis (LDA) model (the “fitcdiscr” function in MATLAB) was trained on normalized spike activity over relative time, achieved by delineating 5 equivalent consecutive intervals of the sequence to classify the position of each interval in the sequence. LDA models were trained on 95% of trials and used to classify the interval position in the remaining 5% of trials. To restrict the analysis to a matched numbers of trials, we combined across sessions and subject neurons recorded during sessions with at least 26 complete sequences of duration shorter than 20s and with a port entry latency shorter than 10s. Subsequently, we restricted the analysis to 26 randomly selected trials. For ensemble decoding, we pooled together separately recorded units. We found the 26-fold cross-validated accuracy for models trained on the activity of randomly selected levels of 10, 50, 100, 500, and 900 units. We performed this analysis 50 times for each level. The same analysis was conducted with the interval positions shuffled to determine the accuracy expected from chance. Two-way ANOVAs were conducted to assess the effects of interval positions and ensemble size, on the decoding accuracy.

For single unit decoding, we performed the analysis described above 26 times in a 26-fold cross-validation approach and averaged performance across all 26 repetitions to find that unit’s accuracy. To account for the variability in decoding accuracy resulting from random selection of trials, the analysis was repeated 20 times. To determine whether individual neurons predicted the position of a given interval above chance, we compared the decoding accuracy of each individual neuron for any time interval with the accuracy of the whole population of neurons after independently shuffling the firing activity of each unit across time intervals.

LDA models were also used to predict whether lever press sequences were complete or incomplete based on the activity during the approach of the lever (1s before LP1, lever approach LA), during the first, third or last interval, during the port approach or during the period of reward consumption. Every time periods were tested separately. Only neurons from sessions with at least 10 complete and 10 incomplete sequences were included in the analysis. The analysis was therefore conducted on the activity of 164 neurons across 20 randomly selected complete and incomplete trials (10 of each) with a 10-fold cross-validation approach. To account for the variability in decoding accuracy resulting from random selection of trials, the analysis was repeated 100 times.

For the analyses described above, neurons with extremely low firing rate (with null firing rate values in more than 75% of trials) were excluded to avoid errors from creating an LDA model on a dataset with too little variance. We assessed whether decoding accuracy significantly departed from chance (shuffled data) using permutation test (Fig S9).

#### Statistical analysis

Data following a normal distribution were subjected to repeated measures analysis of variance. The Kruskal-Wallis test was used when normality assumption was violated. Mean z-scores were compared across complete and incomplete sequences using 2-tailed student t-tests. All analyses were conducted on MATLAB (MathWorks).

### Histology

Electrode sites were labeled by passing a DC current through each electrode, under deep anesthesia with pentobarbital. All rats were perfused intracardially with 1M PBS followed by 4% paraformaldehyde. Brains were extracted, post-fixed in 4% paraformaldehyde for 4–24hr, and transferred in 20% sucrose for >48 h for cryo-protection. To verify the electrodes placement, the brains were sectioned at 50 µm on a cryostat and slices were stained with cresyl violet and analyzed using light microscopy.

## Acknowledgments

This work was supported by the National Institute for Health Research (R01DA035943 to P.H.J), the National Institute on Alcohol Abuse and Alcoholism (R01AA026306 to P.H.J), and the Peter and Traudl Engelhorn foundation (to Y.V.)

## Author Contributions

Conceptualization, Y.V. and P.H.J.; Methodology, Y.V., P.H.J., and D.J.O.; Formal analysis, Y.V., D.J.O and P.H.J.; Investigation, Y.V.; Data curation, Y.V.; Writing – Original Draft, Y.V., D.J.O. and P.H.J.; Writing – Review & Editing, Y.V., D.J.O and P.H.J.; Visualization, Y.V.; Funding Acquisition, P.H.J.; Resources, P.H.J.; Supervision, P.H.J.

## Declaration of Interests

The authors declare no competing interests.

## Notes

### Competing Interest Statement

The authors have declared no competing interest.

## References

Akhlaghpour, H., Wiskerke, J., Choi, J.Y., Taliaferro, J.P., Au, J., and Witten, I.B. (2016). Dissociated sequential activity and stimulus encoding in the dorsomedial striatum during spatial working memory. Elife 5, 1–20.

Apicella, P., Scarnati, E., Ljungberg, T., and Schultz, W. (1992). Neuronal Activity in Monkey Striatum Related to the Expectation of Predictable Environmental. J. Neurophysiol. 68, 945–960.

Bakhurin, K.I., Goudar, V., Shobe, J.L., Claar, L.D., Buonomano, D. V., and Masmanidis, S.C. (2017). Differential encoding of time by prefrontal and striatal network dynamics. J. Neurosci. 37, 854–870.

Balleine, B.W., and O’Doherty, J.P. (2010). Human and rodent homologies in action control: corticostriatal determinants of goal-directed and habitual action. Neuropsychopharmacology 35, 48– 69.

Balleine, B.W., Liljeholm, M., and Ostlund, S.B. (2009). The integrative function of the basal ganglia in instrumental conditioning. Behav. Brain Res. 199, 43–52.

Barnes, T.D., Kubota, Y., Hu, D., Jin, D.Z., and Graybiel, A.M. (2005). Activity of striatal neurons reflects dynamic encoding and recoding of procedural memories. Nature 437, 1158–1161.

Collins, A.L., Greenfield, V.Y., Bye, J.K., Linker, K.E., Wang, A.S., and Wassum, K.M. (2016). Dynamic mesolimbic dopamine signaling during action sequence learning and expectation violation. Sci. Rep. 6, 1–15.

Corbit, L.H., and Janak, P.H. (2010). Posterior dorsomedial striatum is critical for both selective instrumental and Pavlovian reward learning. Eur. J. Neurosci. 31, 1312–1321.

Dalley, J.W., and Robbins, T.W. (2017). Fractionating impulsivity: neuropsychiatric implications. Nat. Rev. Neurosci. 18, 158–171.

Dalley, J.W., Everitt, B.J., and Robbins, T.W. (2011). Impulsivity, compulsivity, and top-down cognitive control. Neuron 69, 680–694.

Dews, P.B. (1978). Studies on Responding Under Fixed-Interval Schedules of Reinforcement: the Scalloped Pattern of the Cumulative Record. J. Exp. Anal. Behav. 29, 67–75.

Dezfouli, A., and Balleine, B.W. (2013). Actions, action sequences and habits: evidence that goal-directed and habitual action control are hierarchically organized. PLoS Comput Biol 9, e1003364.

Dezfouli, A., Lingawi, N.W., and Balleine, B.W. (2014). Habits as action sequences: hierarchical action control and changes in outcome value. Philos Trans R Soc L. B Biol Sci 369.

Dickinson, A. (1989). Expectancy theory in animal conditioning. In Contemporary Learning Theories, S.B. Klein, and R.R. Mowrer, eds. (Psychology Press), p.

Dickinson, A., and Balleine, B. (1994). Motivational Control of Instrumental Action. Anim. Learn. Behav. 22, 1–18.

Donnelly, N.A., Paulsen, O., Robbins, T.W., and Dalley, J.W. (2015). Ramping single unit activity in the medial prefrontal cortex and ventral striatum reflects the onset of waiting but not imminent impulsive actions. Eur. J. Neurosci. 41, 1524–1537.

Emmons, E.B., De Corte, B.J., Kim, Y., Parker, K.L., Matell, M.S., and Narayanan, N.S. (2017). Rodent Medial Frontal Control of Temporal Processing in the Dorsomedial Striatum. J. Neurosci. 37, 8718–8733.

Gouvêa, T.S., Monteiro, T., Motiwala, A., Soares, S., Machens, C., and Paton, J.J. (2015). Striatal dynamics explain duration judgments. Elife 4, 1–14.

Graybiel, A.M., and Grafton, S.T. (2015). The Striatum: Where Skills and Habits Meet. Cold Spring Harb. Perspect. Biol. 7, a021691.

Hart, G., Bradfield, L.A., Fok, S.Y., Chieng, B., and Balleine, B.W. (2018). The Bilateral Prefronto-striatal Pathway Is Necessary for Learning New Goal-Directed Actions. Curr. Biol. 28, 2218–2229.e7.

Hikosaka, O., Sakamoto, M., and Usui, S. (1989). Functional Properties of Monkey Caudate Neurons III. Activities Related to Expectation of Target and Reward. J. Neurophysiol. 61, 814–832.

Howe, M.W., Tierney, P.L., Sandberg, S.G., Phillips, P.E.M., and Graybiel, A.M. (2013). Prolonged dopamine signalling in striatum signals proximity and value of distant rewards. Nature 500, 575–579.

Ito, M., and Doya, K. (2015). Distinct Neural Representation in the Dorsolateral, Dorsomedial, and Ventral Parts of the Striatum during Fixed- and Free-Choice Tasks. J. Neurosci. 35, 3499–3514.

Jin, X., and Costa, R. (2015). Shaping action sequences in basal ganglia circuits. Curr Opin Neurobiol 33, 188–196.

Jin, X., and Costa, R.M. (2010). Start/stop signals emerge in nigrostriatal circuits during sequence learning. Nature 466, 457–462.

Jin, X., Tecuapetla, F., and Costa, R.M. (2014). Basal ganglia subcircuits distinctively encode the parsing and concatenation of action sequences. Nat Neurosci 17, 423–430.

Jog, M.S., Kubota, Y., Connolly, C.I., Hillegaart, V., and Graybiel, a M. (1999). Building neural representations of habits. Science 286, 1745–1749.

Kim, H.R., Malik, A.N., Mikhael, J.G., Bech, P., Tsutsui-Kimura, I., Sun, F., Zhang, Y., Li, Y., Watabe-Uchida, M., Gershman, S.J., et al (2020). A Unified Framework for Dopamine Signals across Timescales. Cell 183, 1600–1616.e25.

Kim, J., Ghim, J., Lee, J.H., and Jung, M.W. (2013). Neural Correlates of Interval Timing in Rodent Prefrontal Cortex. 33, 13834–13847.

London, T.D., Licholai, J.A., Szczot, I., Ali, M.A., Le Blanc, K.H., Fobbs, W.C., and Kravitz, A. V. (2018). Coordinated ramping of dorsal striatal pathways preceding food approach and consumption. J. Neurosci. 38, 3547–3558.

Ma, L., Hyman, J.M., Phillips, A.G., and Seamans, J.K. (2014a). Tracking progress toward a goal in corticostriatal ensembles. J. Neurosci. 34, 2244–2253.

Ma, L., Hyman, J.M., Lindsay, A.J., Phillips, A.G., and Seamans, J.K. (2014b). Differences in the emergent coding properties of cortical and striatal ensembles. Nat. Neurosci. 17, 1100–1106.

Martiros, N., Burgess, A.A., and Graybiel, A.M. (2018). Inversely Active Striatal Projection Neurons and Interneurons Selectively Delimit Useful Behavioral Sequences. Curr. Biol. 28, 560–573.e5.

Matell, M.S., and Meck, W.H. (2000). Neuropsychological mechanisms of interval timing behavior. BioEssays 22, 94–103.

Matell, M.S., Meck, W.H., and Nicolelis, M.A.L. (2003). Interval timing and the encoding of signal duration by ensembles of cortical and striatal neurons. Behav. Neurosci. 117, 760–773.

McGeorge, A.J., and Faull, R.L.M. (1989). The organization of the projection from the cerebral cortex to the striatum in the rat. Neuroscience 29, 503–537.

Mello, G.B.M., Soares, S., and Paton, J.J. (2015). A scalable population code for time in the striatum. Curr. Biol. 25, 1113–1122.

Mink, J.W. (1996). THE BASAL GANGLIA?: FOCUSED SELECTION AND INHIBITION OF COMPETING MOTOR PROGRAMS. Prog. Neurobiol. 50, 381–425.

Narayanan, N.S. (2016). Ramping activity is a cortical mechanism of temporal control of action. Curr. Opi 8, 226–230.

Narayanan, N.S., and Laubach, M. (2006). Delay Activity in Rodent Frontal Cortex During a Simple Reaction Time Task. J. Neurophysiol. 101, 2859–2871.

Parker, K.L., Lamichhane, D., Caetano, M.S., and Narayanan, N.S. (2013). Executive dysfunction in Parkinson’s disease and timing deficits. Front. Integr. Neurosci. 7, 1–9.

Parker, X.K.L., Chen, X.K., Kingyon, J.R., Cavanagh, X.J.F., and Narayanan, N.S. (2014). D 1 - Dependent 4 Hz Oscillations and Ramping Activity in Rodent Medial Frontal Cortex during Interval Timing. 34, 16774–16783.

Rivalan, M., Gregoire, S., and Dellu-Hagedorn, F. (2007). Reduction of impulsivity with amphetamine in an appetitive fixed consecutive number schedule with cue for optimal performance in rats. Psychopharmacol. 192, 171–182.

Robbe, D. (2018). To move or to sense? Incorporating somatosensory representation into striatal functions. Curr. Opin. Neurobiol. 52, 123–130.

Rueda-Orozco, P.E., and Robbe, D. (2015). The striatum multiplexes contextual and kinematic information to constrain motor habits execution. Nat. Neurosci. 18, 453–460.

Sales-carbonell, C., Taouali, W., Khalki, L., Pasquet, M.O., Petit, L.F., Moreau, T., Rueda-orozco, P.E., and Robbe, D. (2018). No Discrete Start / Stop Signals in the Dorsal Striatum of Mice Performing a Learned Action Article No Discrete Start / Stop Signals in the Dorsal Striatum of Mice Performing a Learned Action. Curr. Biol. 28, 3044–3055.e5.

Schmitzer-Torbert, N.C., and Redish, A.D. (2008). Task-dependent encoding of space and events by striatal neurons is dependent on neural subtype. Neuroscience 153, 349–360.

Smith, K.S., and Graybiel, A.M. (2016). Habit formation. Dialogues Clin Neurosci 18, 33–43.

Stalnaker, X.T.A., Berg, B., Aujla, X.N., and Schoenbaum, X.G. (2016). Cholinergic Interneurons Use Orbitofrontal Input to Track Beliefs about Current State. J. Neurosci. 36, 6242–6257.

Thorn, C.A., Atallah, H., Howe, M., and Graybiel, A.M. (2010). Differential Dynamics of Activity Changes in Dorsolateral and Dorsomedial Striatal Loops during Learning. Neuron 66, 781–795.

Vandaele, Y., Noe, E., Cador, M., Dellu-Hagedorn, F., and Caille, S. (2018). Attentional capacities prior to drug exposure predict motivation to self-administer nicotine. Psychopharmacology (Berl). 235.

Vandaele, Y., Mahajan, N.R., Ottenheimer, D.J., Richard, J.M., Mysore, S.P., and Janak, P.H. (2019). Distinct recruitment of dorsomedial and dorsolateral striatum erodes with extended training. Elife 8, 1– 29.

Wassum, K.M., Ostlund, S.B., and Maidment, N.T. (2012). Phasic mesolimbic dopamine signaling precedes and predicts performance of a self-initiated action sequence task. Biol. Psychiatry 71, 846– 854.

Wearden, J.H., and Lejeune, H. (2006). “The stone which the builders rejected…”: Delay of reinforcement and response rate on fixed-interval and related schedules. Behav. Processes 71, 77– 87.

Xu, M., Zhang, S.Y., Dan, Y., and Poo, M.M. (2014). Representation of interval timing by temporally scalable firing patterns in rat prefrontal cortex. Proc. Natl. Acad. Sci. U. S. A. 111, 480–485.

Yin, H.H., Ostlund, S.B., Knowlton, B.J., and Balleine, B.W. (2005). The role of the dorsomedial striatum in instrumental conditioning. Eur. J. Neurosci. 22, 513–523.

