## Supplemental figures for "Dorsomedial striatal activity tracks completion of behavioral sequences"

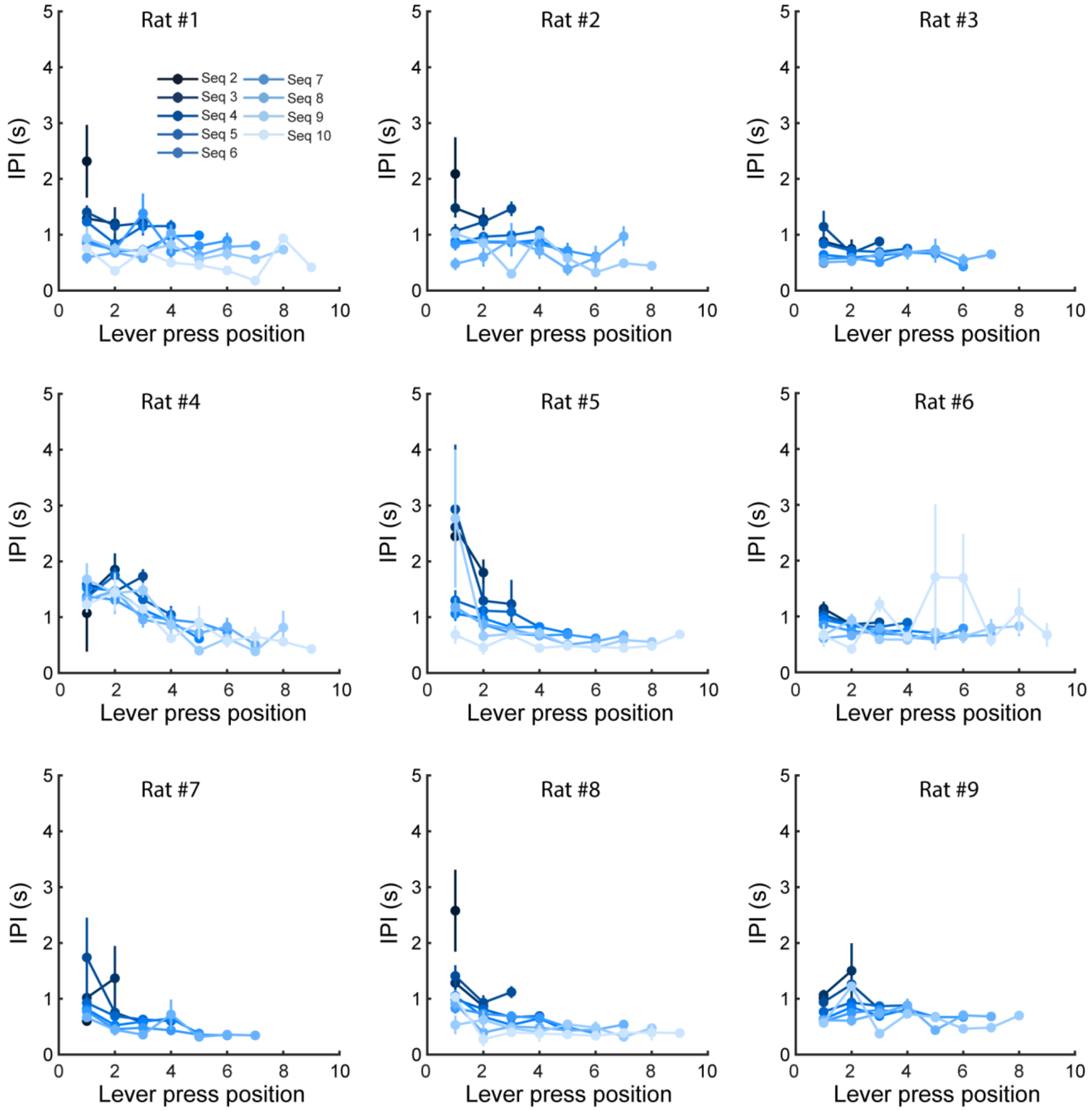


Figure S1: Variations in inter-press intervals within and across sequences in individual rats. Mean inter-press intervals (±SEM) as a function of sequence length and across lever press positions in each individual rat. Refers to Figure 1

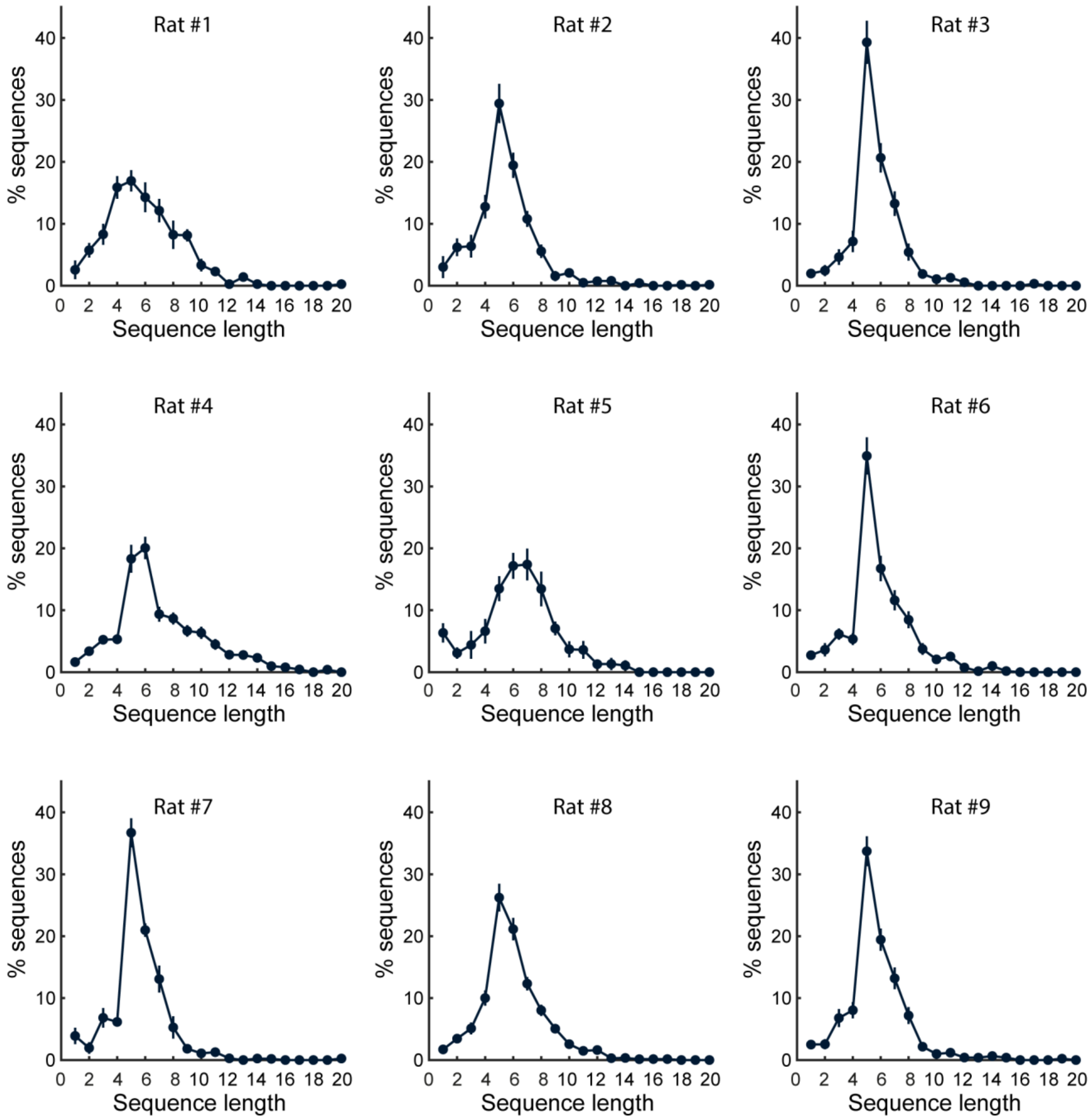


Figure S2: inter-individual variability in sequence length distributions. Mean distribution (±SEM) of sequence length across all analyzed FS5 sessions in each individual rat. Refers to Figure 1

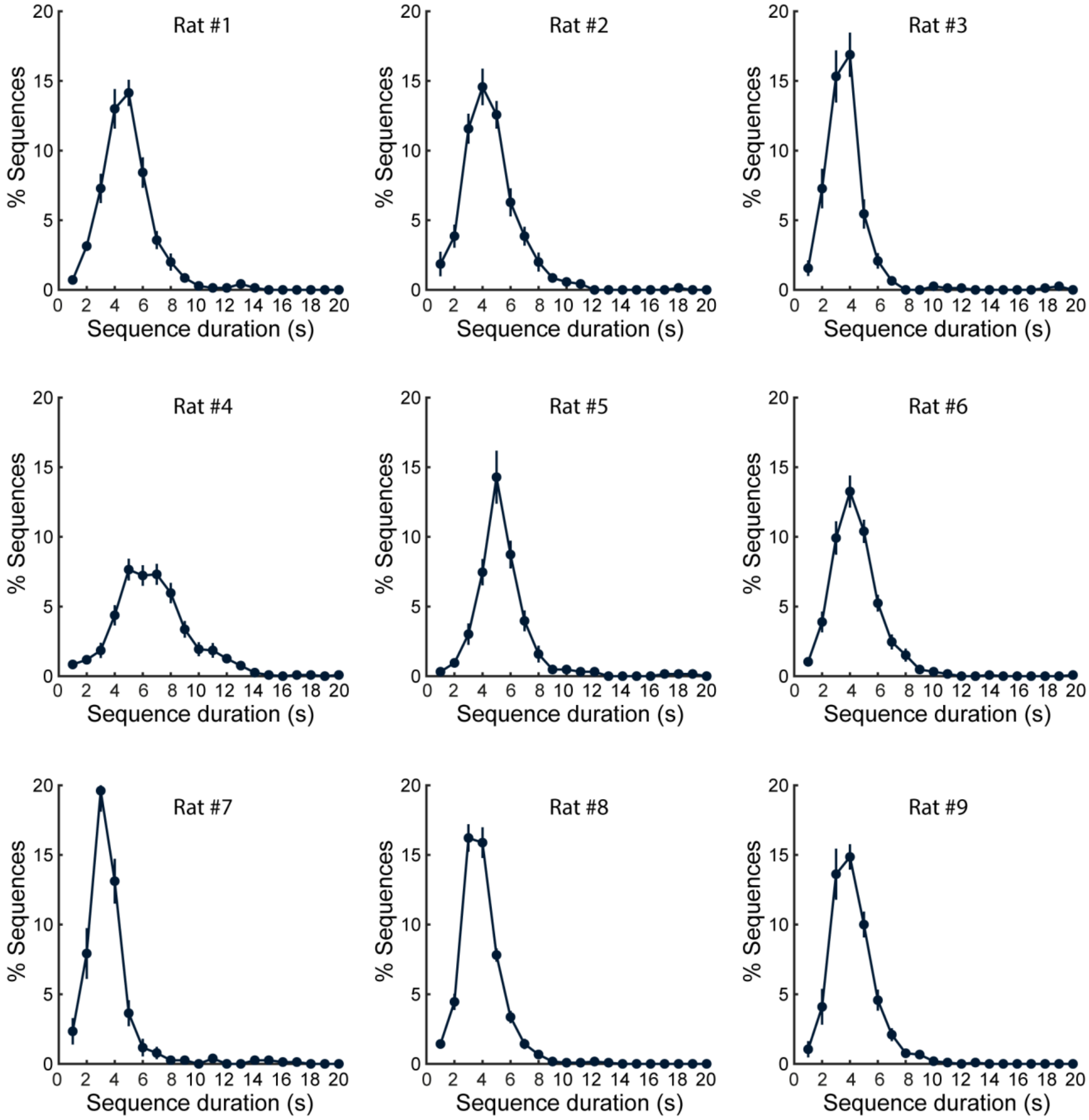


Figure S3: inter-individual variability in sequence duration distributions. Mean distribution (±SEM) of sequence duration across all analyzed FS5 sessions in each individual rat. Refers to Figure 1

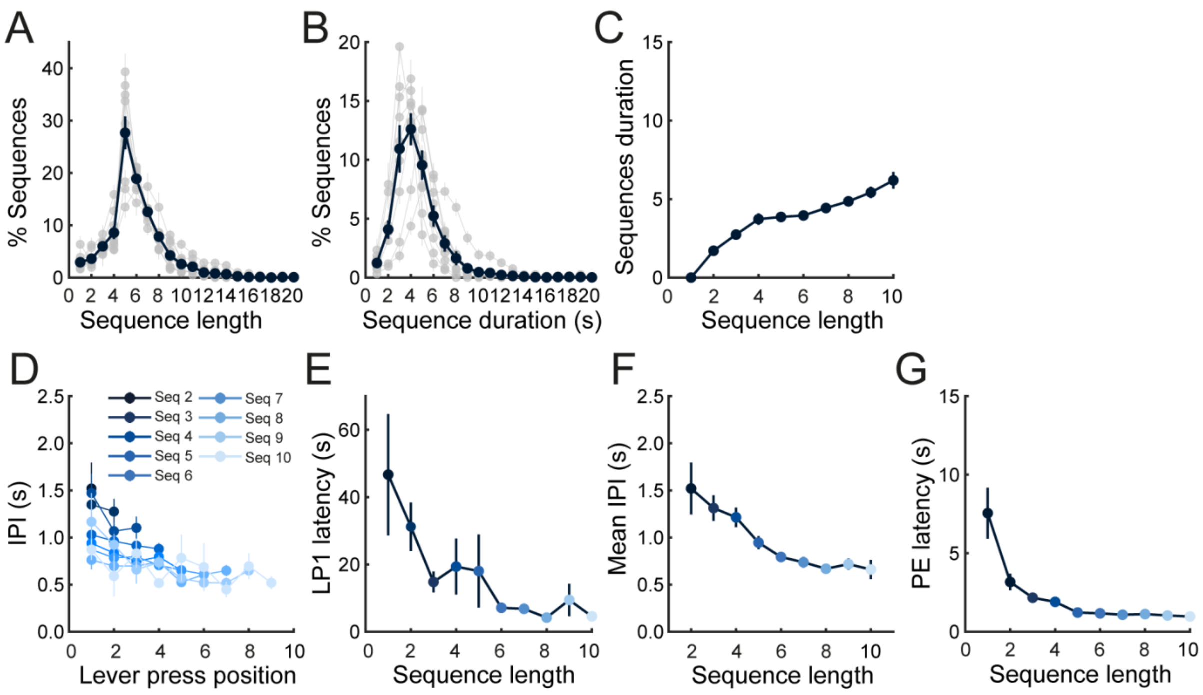


Figure S4: Analysis of behavior combined across sessions and averaged across rats. A-B. Mean distribution of sequence length (A) and duration (B). Gray plots represent individual rats. C. Mean sequence duration as a function of sequence length. D. Mean inter-press intervals (IPI) as a function of sequence length and across lever press position. E-G. Mean LP1 latency (E), IPI (F) and PE latency (G) as a function of sequence length. Data represent the mean (±SEM). Refers to Figure 1

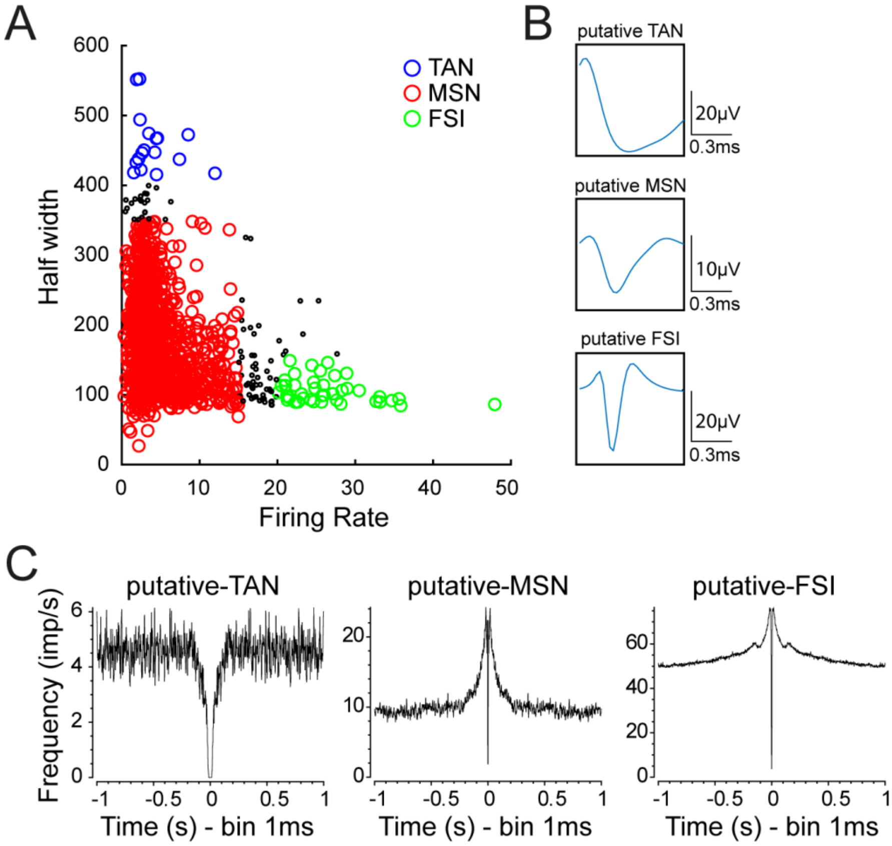


Figure S5: Identification of medium spiny neurons. A. Scatter plot of firing rate and half-valley width allowing separation of tonically active neurons (TAN), medium spiny neurons (MSN) and fast spiking interneurons (FSI). B. Example of the average waveform of putative-TAN, -MSN, and -FSI. C. Example of autocorrelograms of putative-TAN, -MSN, and -FSI. Refers to Figure 2

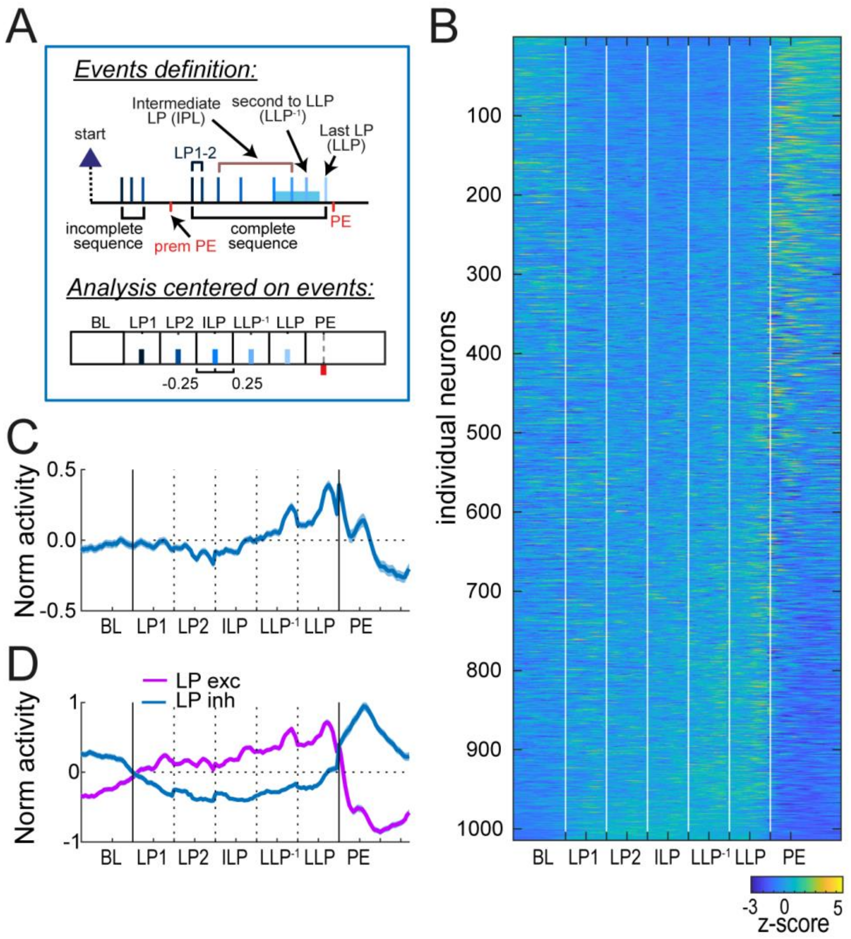


Figure S6: Characterization of DMS activity with an event-centered approach. A. Diagram of task events and analysis. B-C. Heatmap (B) and average z-score (±SEM) (C) of MSNs. D. Average z-score (±SEM) of MSNs excited or inhibited during lever presses. Refers to Figure 2.

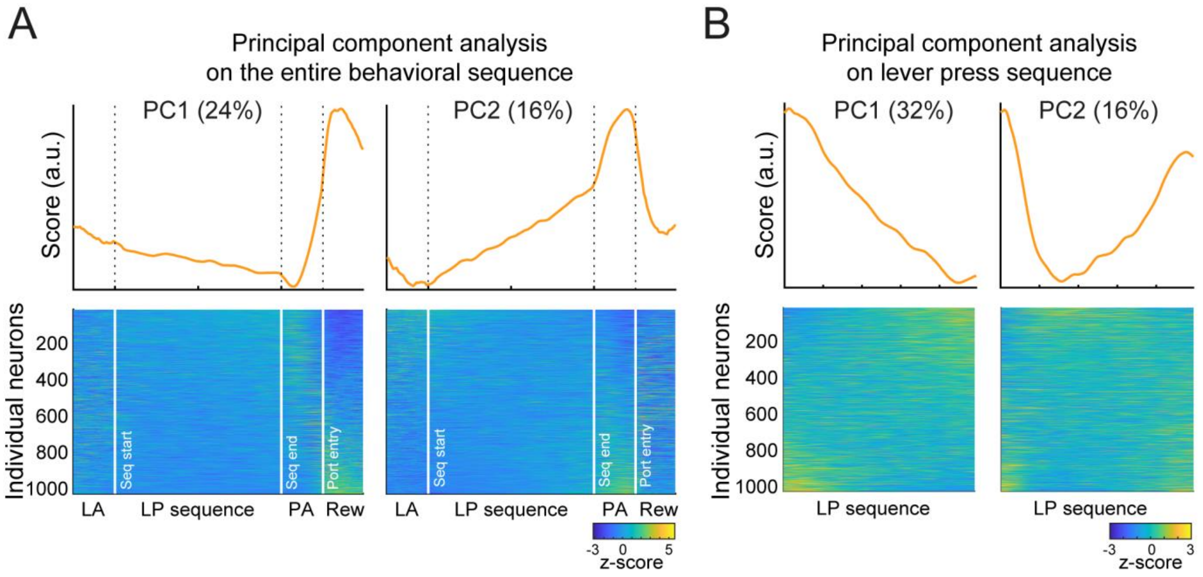


Figure S7: Principal Component Analysis on the entire behavioral sequence (A) or restricted to lever presses (B). A. Eigenvector values of PC1 and PC2 (top) and heatmaps (bottom) of MSNs sorted based on their coefficient for PC1 and PC2 (bottom) with a PCA on the entire behavioral sequence. B. Eigenvector values of PC1 and PC2 (top) and heatmaps (bottom) of MSNs sorted based on their coefficient for PC1 and PC2 (bottom) with a PCA restricted to the lever presses. The % variance explained by each component is indicated. Refers to Figure 2.

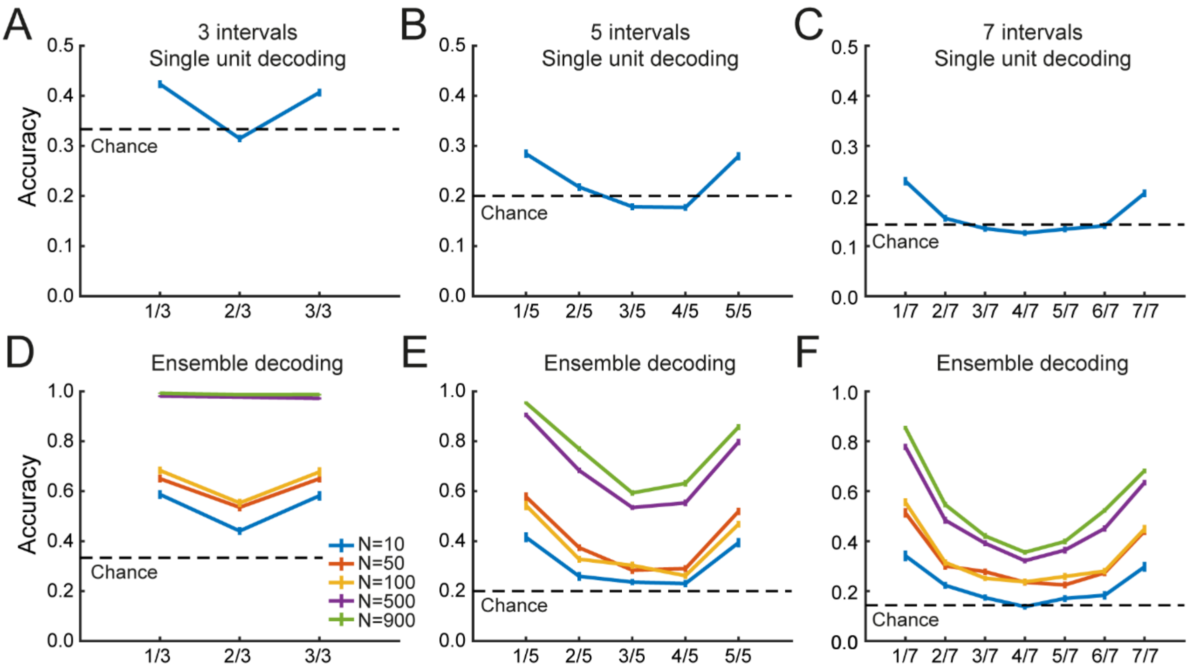


Figure S8: DMS activity pattern tracks progress across time intervals of the behavioral sequence. A-C. Mean single-unit decoding accuracy (±SEM) across time intervals of sequences subdivided in 3 (A), 5 (B) and 7 (C) equivalently-sized consecutive intervals. D-F. Mean decoding accuracy (±SEM) as a function of pseudo-ensemble size and across time intervals of sequences subdivided in 3 (D), 5 (E) and 7 (F) equivalently-sized consecutive intervals. Refers to Figure 3

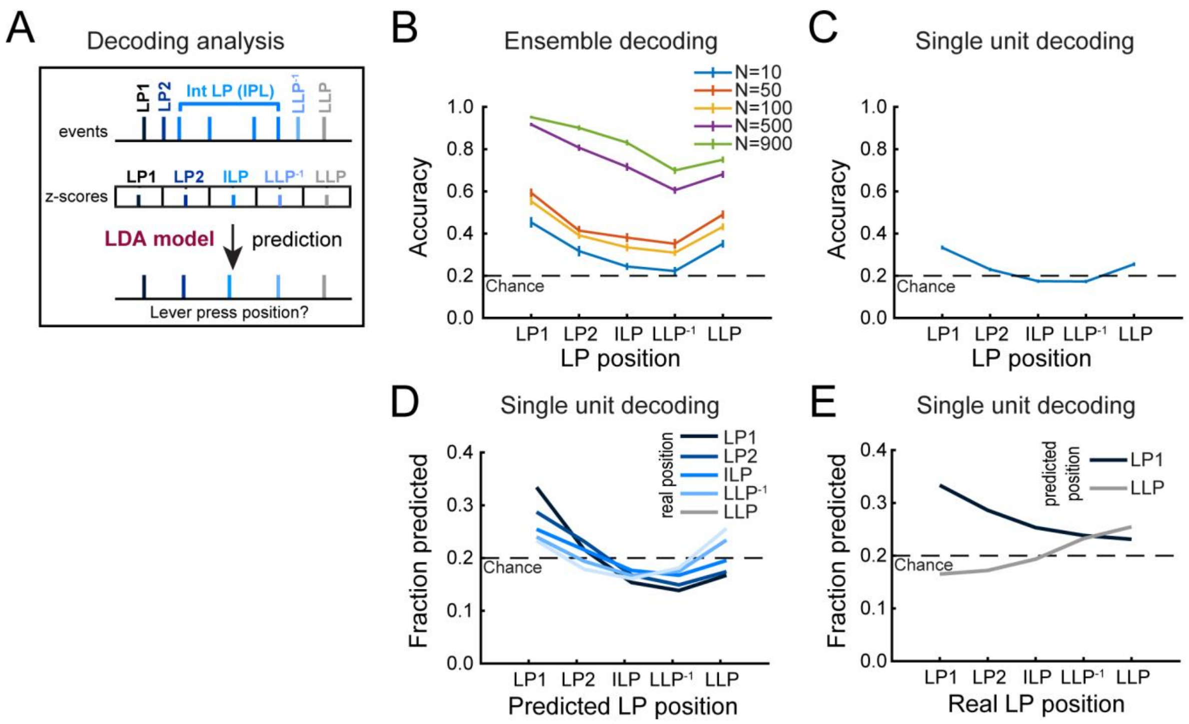


Figure S9: Progress in the lever press sequence is encoded in DMS activity pattern. A. Diagram of the decoding analysis with an event-centered approach. B. Mean decoding accuracy (±SEM) across lever presses and as a function of pseudo-ensemble size. C. Mean single-unit decoding accuracy (±SEM) across lever presses. D. Fraction of predicted lever press position as a function of real lever press position. E. Fraction of lever press predicted as the first and the last press as a function of real lever press position. Refers to Figure 3

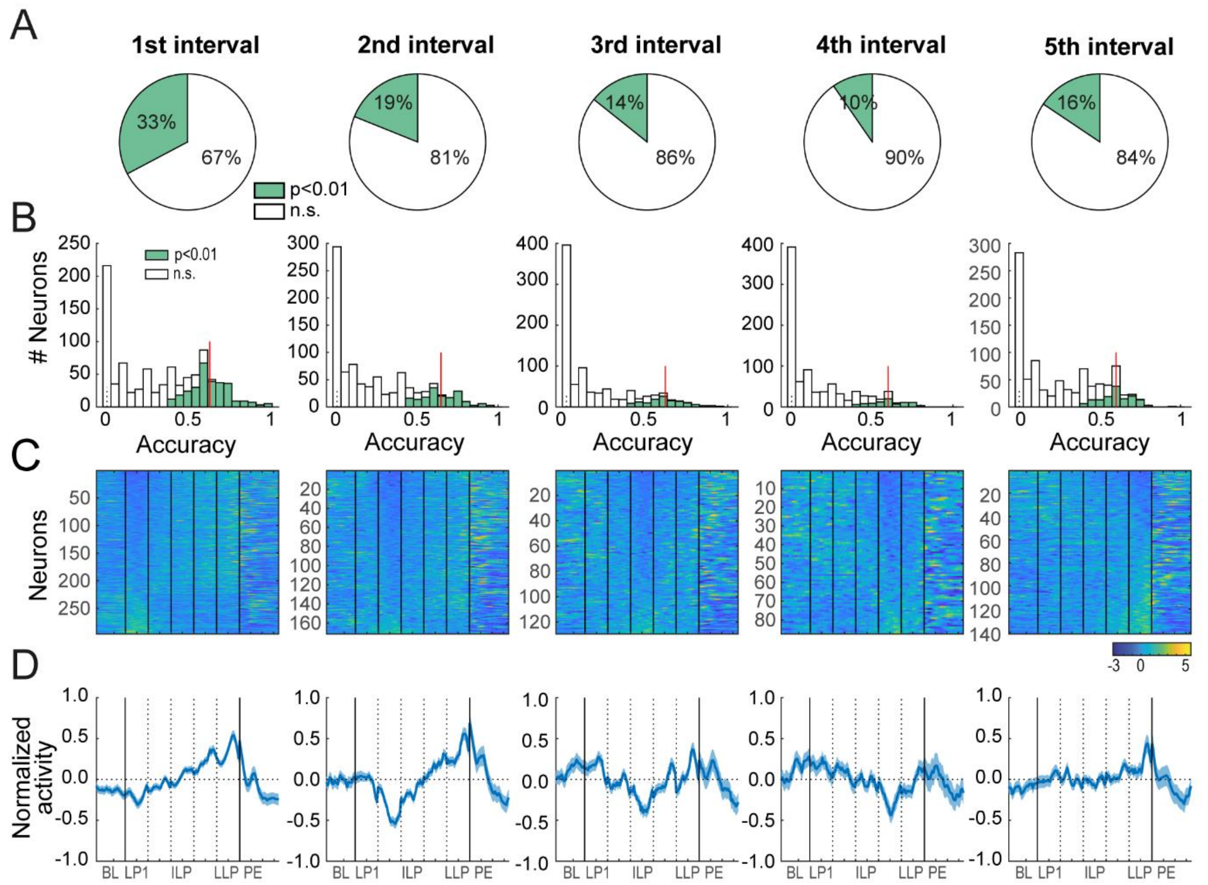


**Figure S10: Single-units individually decode the location of few lever press events**. **A.** Proportion of individual neurons that best predicted the position of each lever press event above chance. **B.** Distribution of decoding accuracy of individual neurons that best predicted each of the lever press events. **C-D.** Heatmaps (C) and average z-score (±SEM) (D) of neurons that best predicted the position of a lever press event above chance, for each lever press. Refers to Figure 4.

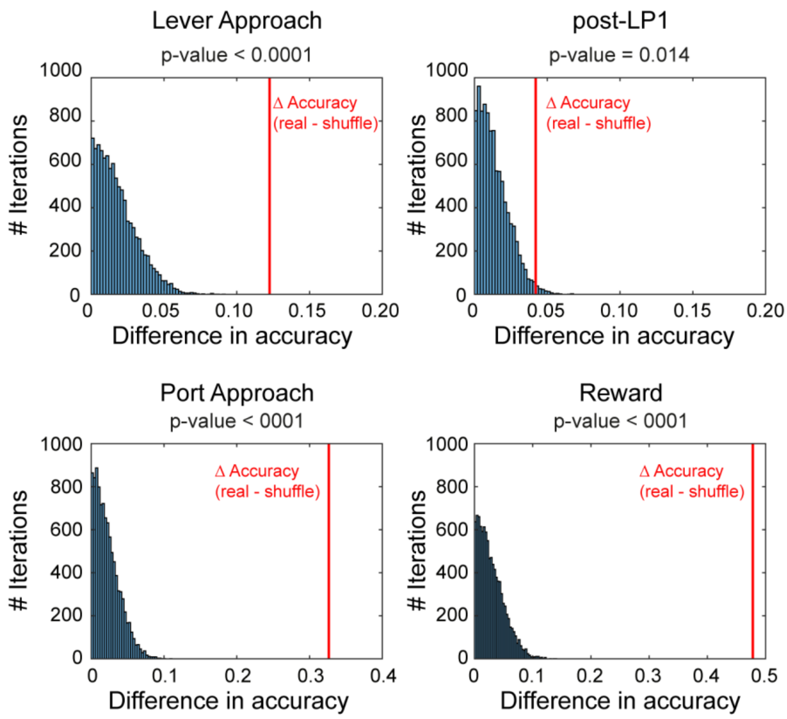


Figure S11: Permutation tests to assess significance of decoding accuracy between complete and incomplete sequences at different time events. We shuffled complete and incomplete sequences across 10000 iterations. In each graph, red vertical lines indicate the difference in accuracy between the real data and the shuffle condition. Refers to Figure 5.

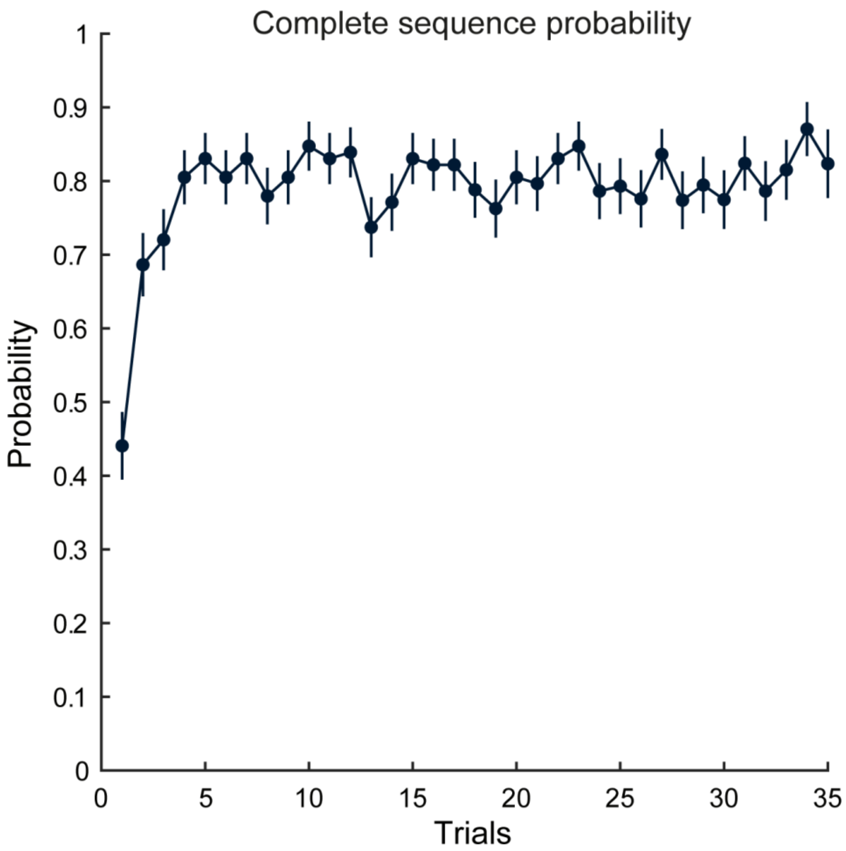


Figure S12: Stable within-sessions repartition of complete and incomplete sequences. Probability of completion of lever press sequence as a function of sequence trials. Refers to Figure 5.
